# Air plasma-activated medium exerts tumor-specific cytotoxicity via the oxidative stress-induced perinuclear mitochondrial clustering

**DOI:** 10.1101/2021.10.31.466636

**Authors:** Manami Suzuki-Karasaki, Takashi Ando, Yushi Ochiai, Kenta Kawahara, Miki Suzuki-Karasaki, Hideki Nakayama, Yoshihiro Suzuki-Karasaki

## Abstract

Intractable cancers such as osteosarcoma (OS) and oral cancer (OC) are highly refractory, recurrent, and metastatic once developed, and their prognosis is still disappointing. Tumor-targeted therapy eliminating cancers effectively and safely is the current clinical choice. Since aggressive tumors have inherent or acquired resistance to multidisciplinary therapies targeting apoptosis, tumor-specific induction of another cell death modality is a promising avenue to meet the goal. Here, we report that a cold atmospheric air plasma-activated medium (APAM) can induce cell death in OS and OC via a unique mitochondrial clustering. This event was named monopolar perinuclear mitochondrial clustering (MPMC) because of the characteristic unipolar mitochondrial perinuclear aggregation. APAM had potent antitumor activity both i*n vitro* and *in vivo*. APAM caused apoptosis, necrotic cell death, and autophagy. APAM contained hydrogen peroxide and increased mitochondrial ROS (mROS), while the antioxidant *N*-acetylcysteine (NAC) prevented cell death. MPMC occurred following mitochondrial fragmentation coinciding with nuclear damages. MPMC was accompanied by the tubulin network remodeling and mitochondrial lipid peroxide (mLPO) accumulation and prevented by NAC and the microtubule inhibitor, Nocodazole. Increased Cardiolipin (CL) oxidation was also seen, and NAC and the peroxy radical scavenger Ferrostatin-1 prevented it. In contrast, in fibroblasts, APAM induced minimal cell death, mROS generation, mLPO accumulation, CL oxidation, and MPMC. These results suggest that MPMC is a tumor-specific cause of cell death via mitochondrial oxidative stress and microtubule-driven mitochondrial motility. MPMC might serve as a promising target for exerting tumor-specific cytotoxicity.

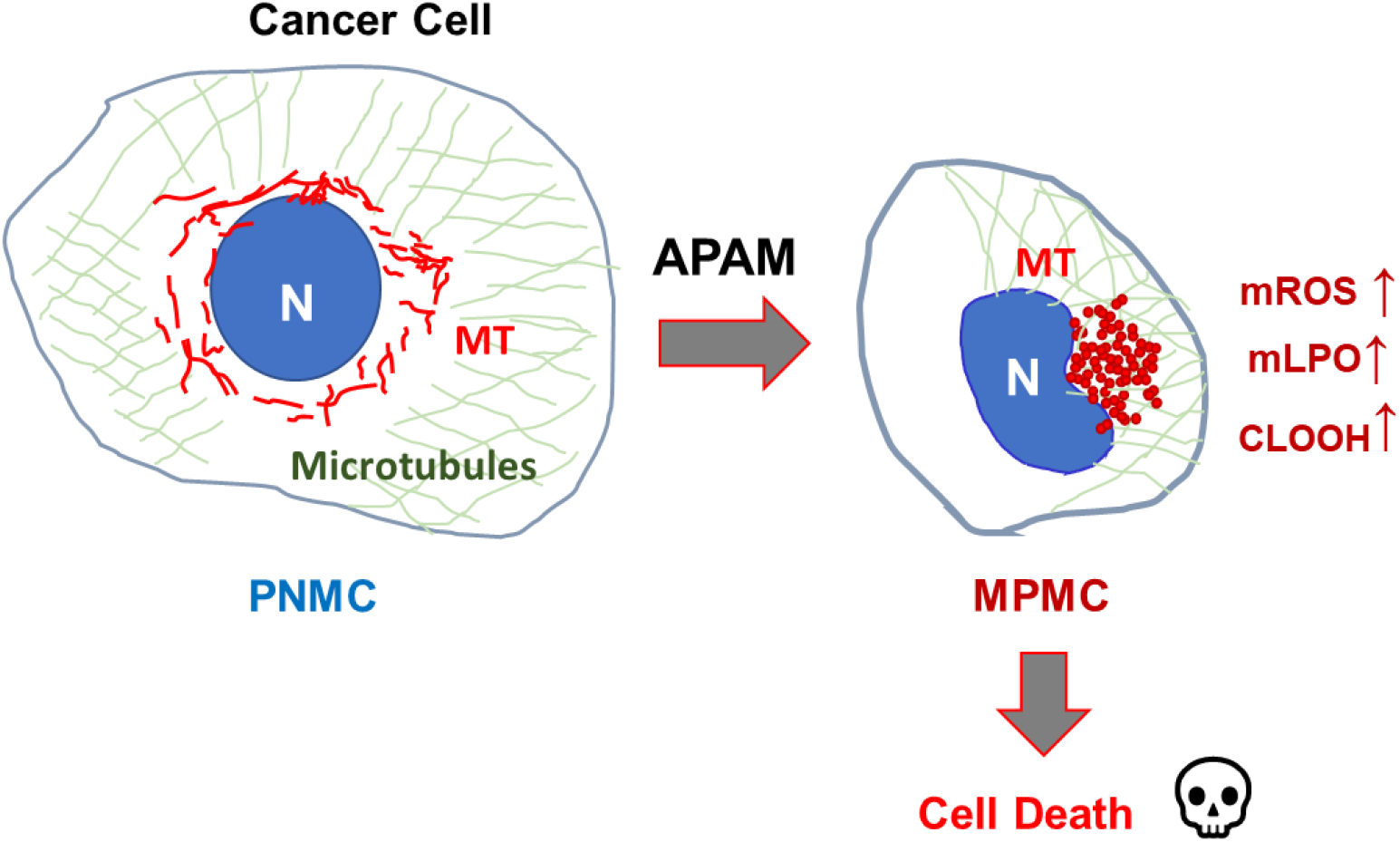
Graphical abstract.

## Introduction

Intractable cancers such as osteosarcoma (OS) and oral cancer (OC) are greatly refractory, recurrent, and metastatic once developed, and their prognosis is still disappointing. Tumor-targeted therapy eliminating cancers effectively and safely is the current clinical choice. The activation of tumor-selective cell death is a promising approach to meet the goal. Apoptosis is the primary modality of cancer cell death caused by diverse chemical and physical stresses and multidisciplinary therapies, including anticancer agents. However, it is evident that these aggressive tumors have cellular machinery protecting them from apoptosis, conferring inherent or acquired resistance (1‒3). An emerging view is that apoptosis is not the sole mode of cancer cell death. Several different ways of cell death, such as autophagy and necroptosis, also occur in response to anticancer agents (4, 5). Thus, activation of nonapoptotic cell death can be a promising alternative approach to treating intractable cancers.

Cold atmospheric plasma (CAP) has recently emerged as a promising tumor-targeting approach. Direct CAP treatment inhibits cell proliferation, migration, and invasion. It triggers different cell death modalities, including apoptosis, necrosis, and autophagy *in vitro* in various cancer cell lines and primary cancerous cells and tissues (6‒10). CAP also reduced cell growth of xenografted tumors *in vivo* while exhibiting minimal toxicity in normal cells and tissues (7, 9). These properties have attracted much attention in cancer treatment. Exposing CAP to various solutions results in plasma-activated liquids (PALs) such as plasma-activated media (PAM). PALs have different chemical and biological effects depending on the physical properties of plasma generated and various experimental parameters. Despite such variation, different types of PAM have antitumor activity against tumor cells both *in vitro* and *in vivo* while sparing normal cells and tissues (11‒16). Helium plasma-activated media (HePAM) and plasma-activated infusion (PAI) activate autophagy and necroptosis in a tumor-selective manner (13, 14, 17). These PALs also have antitumor activity against OS both *in vitro* and *in vivo* with minimal side effects. Additionally, they can be easily administered systematically or locally to deep tissues by infusion and endoscope. Thus, they will promise a potent and safe approach for cancer treatment.

Mitochondria are highly dynamic organelles, constantly changing their size, shape, and location to cope with the energy demands (18, 19). A growing body of evidence indicates that mitochondrial network dynamics play a critical role in regulating functions and survival in non-transformed and transformed cells. The dynamics are regulated by the delicate balance between fission and fusion of the mitochondrial membrane. Dynamin-related proteins (Drps) with GTPase activity such as Drp1, optic atrophy 1 (OPA1), and mitofusin 1/2 (Mfn1/2) act in concert to regulate the dynamics; Drp1 regulates fission while OPA1 and Mfn1/2 control fusion and cristae organization. A defect in either process causes severe mitochondrial and cellular dysfunctions (20, 21). Mitochondrial fission facilitates eliminating damaged mitochondria through mitophagy-mediated disruption. Therefore, its deficiency leads to a hyperfused mitochondrial network and compromised mitochondrial quality control (22). On the other hand, mitochondrial fusion facilitates the exchange of mitochondrial DNA and metabolites required for mitochondrial function. Accordingly, its failure leads to mitochondrial fragmentation, reduced mitochondrial DNA, growth, mitochondrial membrane potential (ΔΨ_m_), and defective respiration (23). Mitochondrial motility and subcellular positioning have recently emerged as another critical regulatory factor in mitochondrial and cellular functions (24, 25). In the perinuclear mitochondrial clustering (PNMC), mitochondria are clustered around perinuclear sites. PNMC has been suggested to maintain the hypoxic status of cancer cells via mitochondrial ROS production and stabilization of hypoxia-inducible factor 1α (HIF-1α), the master transcription factor regulating hypoxic signaling (26–29).

We previously reported that tumor necrosis factor-related apoptosis-inducing ligand (TRAIL) induced mitochondrial fragmentation by increasing Drp1 phosphorylation at Ser 616. The Drp1 inhibitor Mdivi-1 and Drp1 gene silencing significantly augmented TRAIL-induced caspase-3 activation, mitochondrial depolarization, and apoptosis (30, 31). These facts indicate that the Drp1-dependent mitochondrial fragmentation is cytoprotective. The enhancement of apoptosis pathways is associated with the shift from mitochondrial fragmentation to clustering, and increased plasma membrane depolarization causes it (31). Strikingly, such a shift occurs in OS, malignant melanoma, and lung cancer, but not normal cells. HePAM (12, 13) and PAI (17) also induce mitochondrial clustering in various cancer cells. They quickly cause mitochondrial fragmentation at low (subtoxic) concentrations while causing mitochondrial clustering at high (toxic) doses. On the other hand, even when applied at high concentrations, they caused substantial fragmentation only in non-transformed cells such as melanocytes and fibroblasts. Together, these facts suggest that cancers are more prone than normal cells to mitochondrial clustering, and this higher susceptibility may lead to tumor-selective cell death induction. Thus, it is worth knowing whether the given antitumor agent can evoke mitochondrial clustering.

Recently, we noticed that air bubbling augmented the antitumor activity of HePAM, suggesting the involvement of air components in action. We used the ambient air instead of helium as a CAP source to enrich the active substances, generating an air plasma-activated medium (APAM). In the present study, we explored the antitumor activity of APAM with a particular interest in its impact on the mitochondrial network and positioning. We found that APAM induced nonapoptotic cell death via a unique death-associated mitochondrial perinuclear clustering in a tumor-selective manner.

## 2. Results

### 2.1. APAM has potent cytotoxicity against OS in *vitro* and *in vivo*

First, we examined whether APAM had antitumor activity *in vitro*. Cells were treated with APAM at different concentrations (7, 12.5, 25, 50% solution) for 72 h and analyzed for cell viability by a WST assay. APAM treatment significantly reduced cell viability in HOS, LM8, and 143B OS cells in a dose-dependent manner (Fig. 1A–C). However, the degree of the reduction varied considerably depending on the cell line. APAM (≤12.5%) caused a substantial effect (≥50% reduction) in high responders, including human HOS cells. In comparison, APAM (≥25%) was necessary for a ≥50% reduction in low responders such as human 143B and murine LM8 cells (Fig. 1A–C). The effect was preceded by cellular and nuclear morphological changes, including appearing in blebbed, nonadherent, round cells with shrunken or fragmented nuclei (see Fig. 3). Next, we examined whether APAM had antitumor activity *in vivo*. The LM8 cells can be transplanted into the host C3H mice to develop solid tumors. We assessed the action in the allograft model under the regimen shown in Supplementary Fig. S1. Subcutaneous inoculation of the cells into mice resulted in rapid tumor growth, reaching about 3000 mm^3^ within five weeks. Intravenous administration of APAM (50%) three times a week resulted in a significant (68.1%) reduction in the tumor size (Fig. 1D). APAM treatment also reduced tumor growth in a xenograft model where human 143B cells were inoculated into nude mice. The tumor developed about 5000 mm^3^ within five weeks after inoculation. APAM administration resulted in a significant (74.2%) reduction in the tumor size (Fig. 1F). There was a minimal difference in mice’s weights between the treated and untreated groups and any symptom of adverse effects (Fig. 1E, G). These results indicate that APAM has potent tumor-targeting anticancer activity against OS *in vitro* and *in vivo*.

**Fig. 1.**
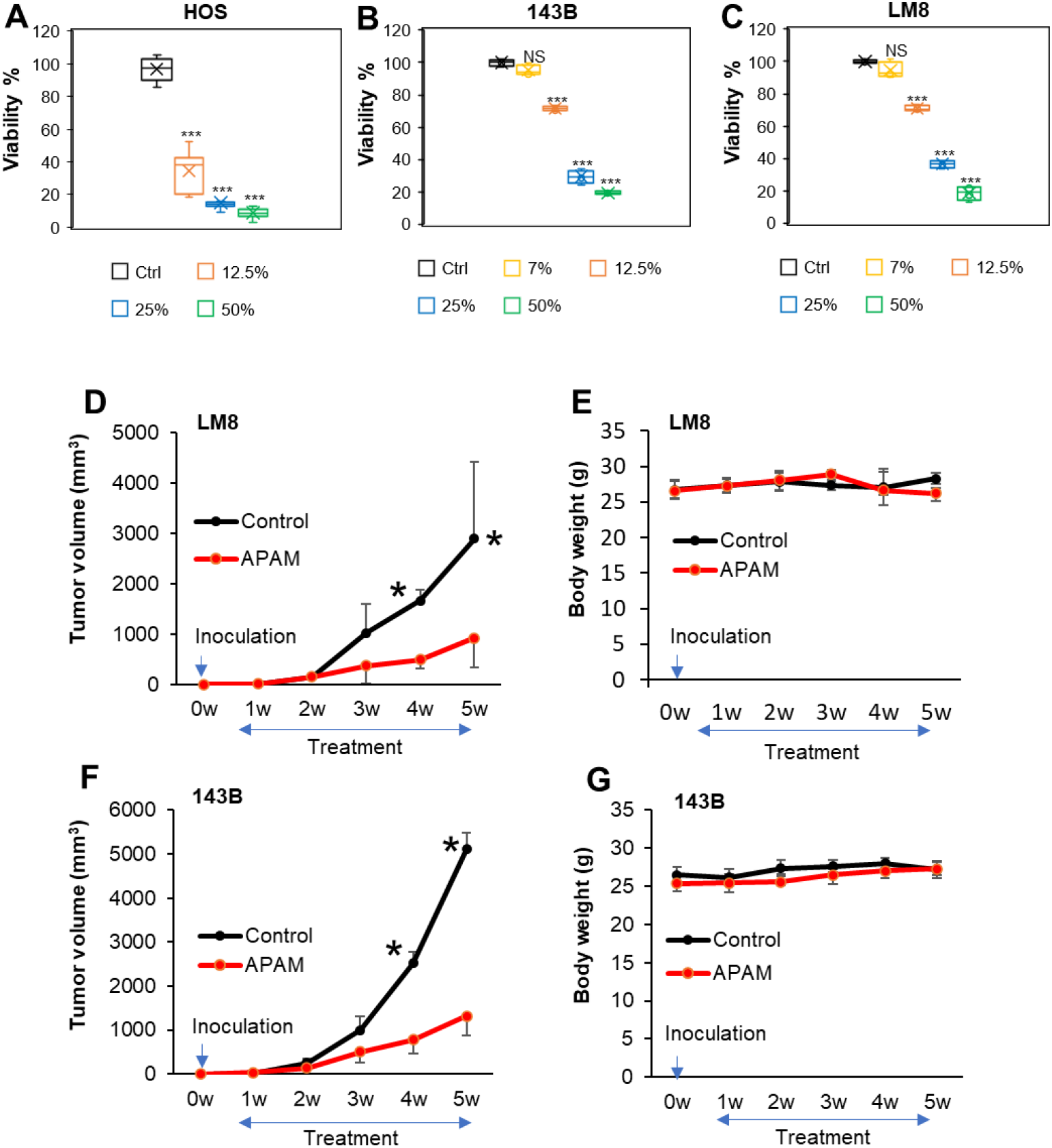
APAM has potent antitumor activity against OS *in vitro* and *in vivo*. A‒C. HOS (A), 143B (B), and LM8 (C) cells (4×10^3^ cells) were treated with the indicated concentrations of APAM for 72 h and analyzed for viability using a WST-8 cell growth assay. Data are the mean ± SD (n = 4‒9). Data were analyzed by one-way analysis of variance followed by Tukey’s post hoc test. \*\*\**P <* 0.001; NS, not significant vs. control treated with vehicle. D‒G. C3H (D, E) and nude mice (F, G) were inoculated with 1×10^6^ each of LM8 (D, E) and 143B cells (F, G), respectively at day 0, and intravenously administered 200 μL of APAM (50%) 3 times per week for 4 weeks from day 7. The sizes of the tumors in the mice (D, F) and mice’ weights (E, G) were measured weekly. Values represent the mean ± SD (n = 6). \**P <* 0.05 vs. the control treated with vehicle.

**Fig. 2.**
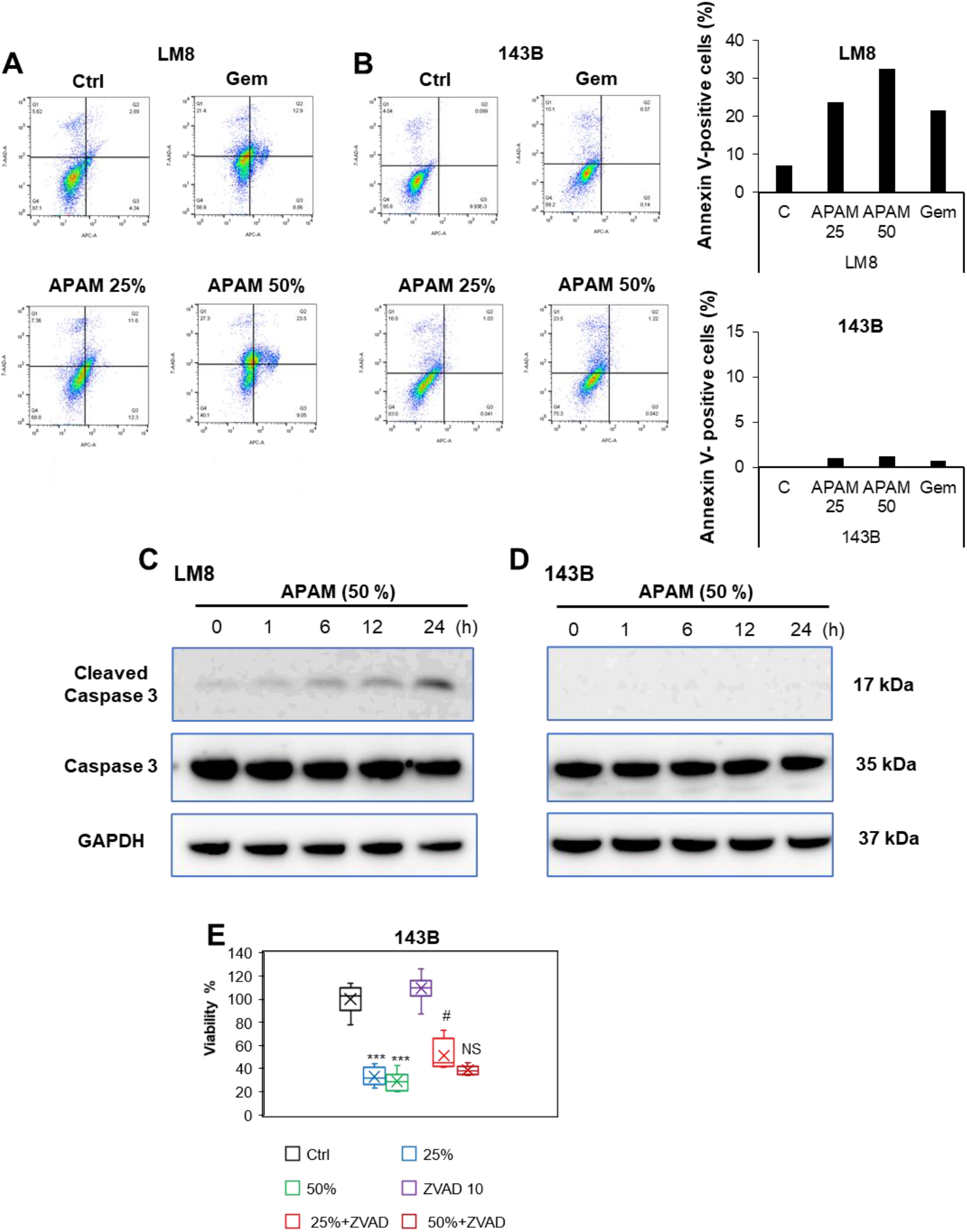
APAM induces apoptotic and nonapoptotic cell death. A, B. LM8 (A) and 143B (B) cells were treated with APAM (25, 50%) for 24 h, stained with Annexin V-APC and 7-AAD, and analyzed by flow cytometry. Gemcitabine (Gem) was used as a positive control for apoptosis induction. The ratios of apoptotic (Annexin-V-positive) cells were shown in the right panels. C, D. LM8 (C) and 143B cells (D) were treated with APAM (50%) for the indicated time and then analyzed for the expression of the full-length and cleaved (active) Caspase-3 by western blotting analysis using specific antibodies. GAPDH was used as the loading control. See Supplementary Fig. S2 for examples of uncropped images and quantification for each antibody. E. 143B cells were pretreated with 10 μM *Z-*VAD-FMK for 1 h and then treated with APAM (25, 50%) for 72 h and analyzed for viability using the WST-8 cell growth assay. Data are the mean ± SD (n = 4‒9). Data were analyzed by one-way analysis of variance followed by Tukey’s post hoc test. \*\*\**P <* 0.001 vs. control treated with vehicle, # *P <* 0.05; NS vs. APAM alone.

**Fig. 3.**
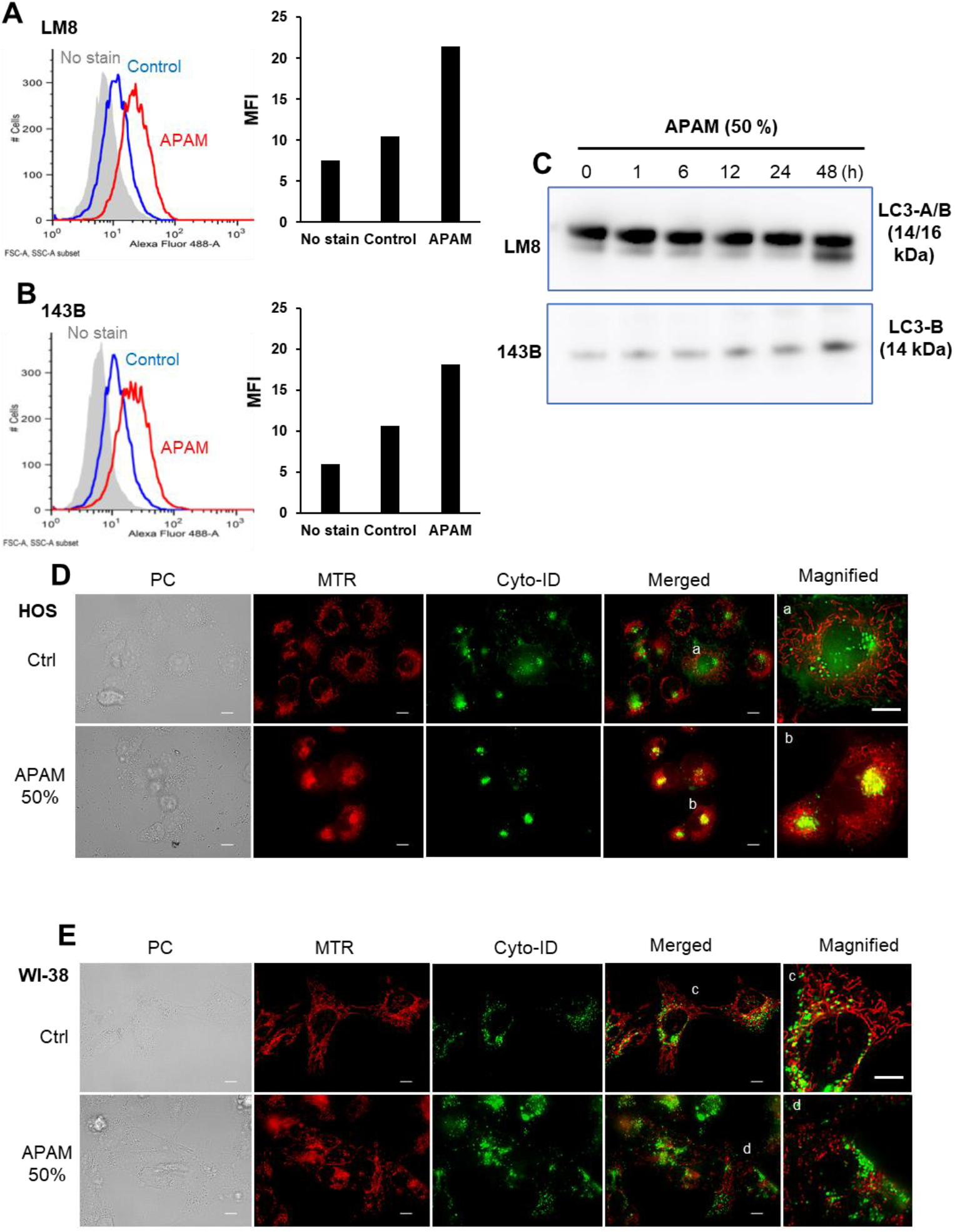
APAM induces autophagic flux and mitophagy. A, B. LM8 (A) and 143B (B) cells were treated with APAM (50%) for 18 h and stained with DAL Green and analyzed by flow cytometry. The right panels show mean fluorescence intensity (MFI). C. LM8 (upper) and 143B cells (lower) were treated with APAM (50%) for the indicated time and analyzed for the expressions of LC3-A/B (upper) and LC3-B (lower), respectively by western blotting analysis using specific antibodies. The loading control was the same as Fig. 2. See Supplementary Fig. S3 for examples of uncropped images and quantification for each antibody. D, E. After treating HOS (D) and WI-38 cells (E) with APAM (50%), mitochondria and autophagosomes were stained with MTR and Cyto-ID, respectively. Images were obtained using a BZ710 Digital Biological Microscope equipped with a 100 × coverslip-corrected oil objective and analyzed using BZ-H3A application software. The right panels (a‒d) show the magnification of a‒d in the merged images. PC, phase contrast. Bar = 10 μm.

### 2.2. APAM can activate apoptosis and nonapoptotic cell death in OS cells

To investigate cell death modality, we analyzed Allophycocyanin (APC)-conjugated Annexin V and 7-aminoactinomycin D (7-AAD) double staining. The cells were treated with APAM (25, 50% solution) for 24 h and analyzed for their Annexin V and 7-AAD double staining in a flow cytometer. Figure 2 shows the results obtained in LM8 and 143B cells. Gemcitabine (Gem) was used as an apoptosis inducer in OS (17). APAM resulted in dose-dependent increases in apoptotic (Annexin V-positive) cells and necrotic (Annexin V-negative) cells in LM8 cells (Fig. 2A). Apoptosis was increased up to a maximum of 30% in response to APAM (50%). On the other hand, APAM caused minimal apoptotic cells while increasing necrotic cells in 143B cells in a dose-dependent manner (Fig. 2B). Western blotting analyses using cleaved caspase-3-specific antibodies confirmed the flow cytometric observations. APAM treatment increased the cleaved form of caspase-3 (17 kDa), a hallmark of apoptosis, over time in LM8 cells (Fig. 2C), but not in 143B cells (Fig. 2D). Additionally, the broad-spectrum caspase inhibitor Z-VAD-FMK could not entirely inhibit APAM (50%)’s cytotoxic activity in143B cells (Fig. 2E). On the other hand, the inhibitor attenuated the effect of APAM (25%). These results indicate that APAM can activate apoptosis and nonapoptotic cell death depending on the cell line and concentration.

### 2.3. APAM induces autophagy and mitophagy in OS cells

We previously reported that a HePAM could increase the autophagic flux and autophagic cell death in OS cells (14). HePAM also can induce mitophagy. In contrast, HePAM activated minimal autophagy, mitophagy, and autophagic cell death in fibroblasts (14). Therefore, we examined whether APAM also induced autophagy and mitophagy. Staining with the autolysosome dye, DAL Green, followed by flow cytometry, revealed that APAM increased DALGreen fluorescences in LM8 and 143B cells (Fig. 3A, B). Moreover, Western blotting analysis using specific antibodies showed that APAM increased the expression of LC3-B, the particular hallmark of autophagosome formation in these cells (Fig. 3C). To determine the onset of mitophagy, we stained mitochondria and autophagosomes with MitoTracker Red CMXRos (MTR) and the autophagosome dye Cyto-ID, respectively. Cyto-ID can detect LC3-B without transfection. We observed substantial diffused and partially clustered Cyto-ID puncta in untreated HOS cells even under nutritional and stress-free conditions, indicating the occurrence of ambient autophagy (Fig. 3D, upper panels). However, the puncta were not overlapped with mitochondria in merged images. APAM increased clustered puncta coinciding with mitochondria in the cells (Fig. 3D, lower panels), indicating increased colocalization of mitochondria and autophagosome. On the other hand, a smaller number of Cyto-ID puncta were observed in fibroblasts, and APAM minimally affected the morphology and localization of the puncta in the cells (Fig. 3E). These results suggest that APAM can induce autophagy and mitophagy in a tumor-selective manner.

### 2.4. ROS plays a vital role in cell death

H_2_O_2_ is found in various PALs, including PAM, and represents the primary mediator of anticancer activity (6–9). Therefore, we examined whether APAM contained H_2_O_2_ by directly quantitating its contents in APAM using Amplex Red. We found that APAM contained < 10 μM of H_2_O_2_ (6.1 ± 2.5 μM, n = 4) under the standard experimental conditions (1 min-exposure/mL medium). Next, we determined whether ROS contributed to APAM’s cytotoxic activity. Cells were pretreated with the antioxidant NAC for 1 h and then treated with APAM (25, 50%) for 72 h. The NAC treatment entirely prevented the decrease in cell viability in HOS cells (Fig. 4A), indicating the critical role of ROS in cell death. To explore the role of ROS further, we examined APAM’s ability to induce intracellular ROS production. APAM significantly increased mitochondrial O_2_^•-^ in the cells, as shown by the increased fluorescence of the mitochondria-localizing O_2_^•-^-reactive probe MitoSOX Red (MitoSOX) (Fig. 4B). These results indicate that ROS plays a vital role in OS cell death caused by APAM.

**Fig. 4.**
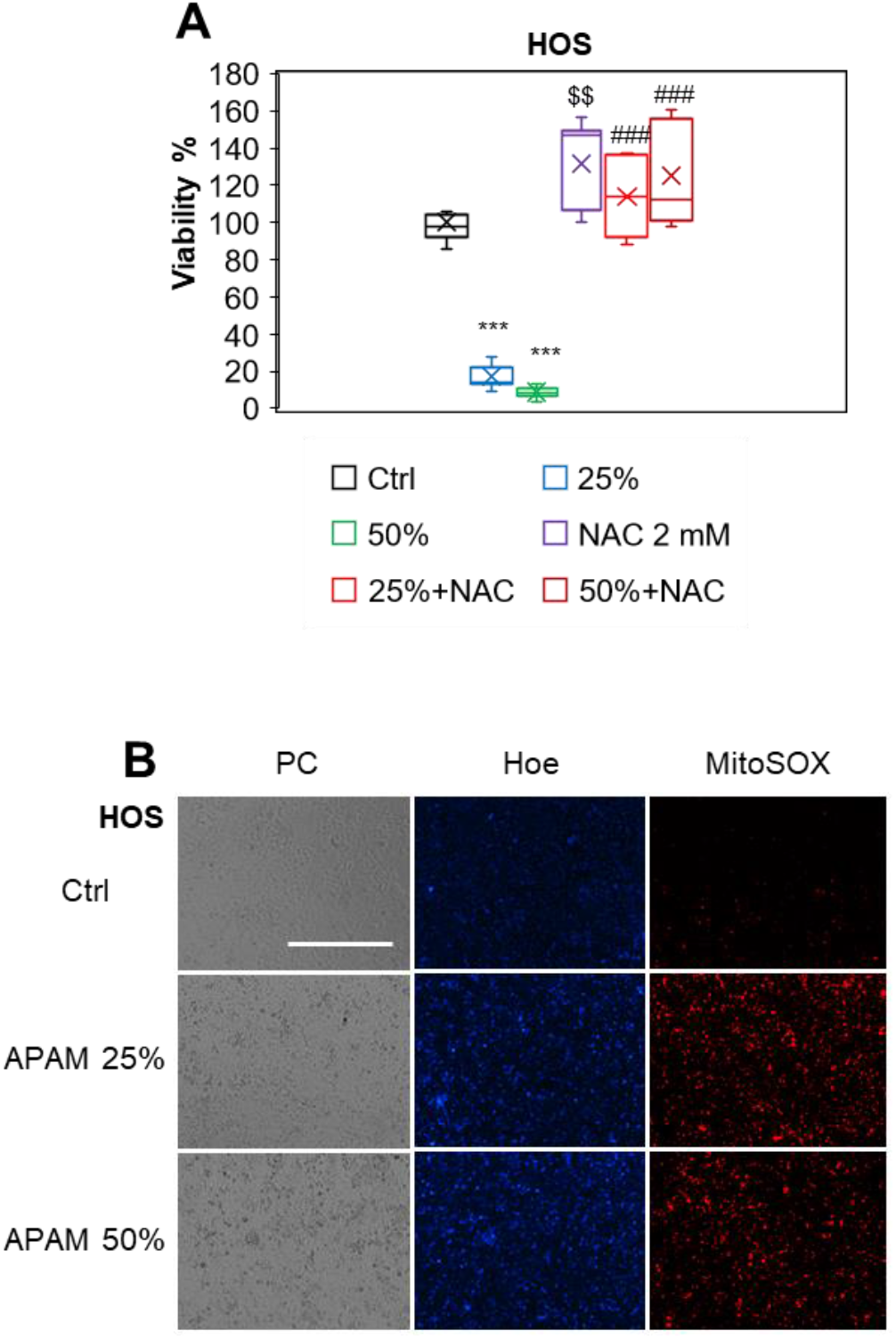
APAM induces cell death in a ROS-dependent manner. A. HOS cells were treated with APAM (25, 50%) or 2 mM NAC alone or in combination for 72 h and analyzed for viability using the WST-8 cell growth assay. Data are the mean ± SD (n = 8 or 9). Data were analyzed by one-way analysis of variance followed by Tukey’s post hoc test. \*\*\**P <* 0.001; $$*P <* 0.01 vs. control treated with vehicle, ###*P <* 0.001 vs. APAM alone. B. HOS cells (1.5×10^4^) were treated with APAM (25, 50%) for 2 h and then stained with 5 μM MitoSOX. The nuclei were counterstained with Hoechst 33342 (Hoe). Images were obtained with EVOS FL Cell Imaging System (Thermo Fisher Scientific) equipped with a 10 × objective and analyzed using the freely available NIH ImageJ software. PC, phase contrast. Bar = 400 μm.

### 2.5. APAM evokes MPMC in a tumor-selective manner

Next, we determined whether APAM affected the mitochondrial network in OS cells. After APAM treatment, the mitochondria in live cells were stained with MitoTracker Red (MTR), and the nuclei were counterstained with Hoechst 33342. The mitochondria in untreated HOS cells had substantial membrane potential and exhibited a reticular network (Fig. 5A, upper panels). Following APAM treatment, mitochondria became fragmented and clustered concomitant with decreased membrane depolarization, as indicated by the dropped MTR signals (Fig. 5A, lower panels). The mitochondrial abnormalities, including the membrane depolarization, were explicitly seen in the round damaged cells (indicated by yellow arrows), indicating that they are death-associated events. Moreover, we noticed that the mitochondrial network collapse was associated with altered mitochondrial subcellular positioning. In untreated cells, the mitochondria are widely distributed through the cytoplasm or all-around or both sides of the perinuclear sites (Fig. 5A, upper panels). On the other hand, following APAM treatment, most mitochondria clustered at one side of the perinuclear sites (Fig. 5A, lower panels). Similar altered mitochondrial morphology and positioning were seen in 143B cells (Fig. 5B). The modified positioning was associated with massive nuclear damages. In contrast, APAM caused only a modest mitochondrial fragmentation with minimal clustering in fibroblasts (Fig. 5C). The quantitative measurement of the mitochondrial occupied area revealed a significant reduction in APAM-treated HOS and 143B cells compared with untreated cells (Fig. 5D, E). On the other hand, no significant decrease was seen in fibroblasts (Fig. 5F). These results indicate that APAM modulates mitochondrial positioning in a tumor-selective manner. Therefore, we named this event monopolar perinuclear mitochondrial clustering (MPMC) and investigated the mechanisms underlying it in more detail.

**Fig. 5.**
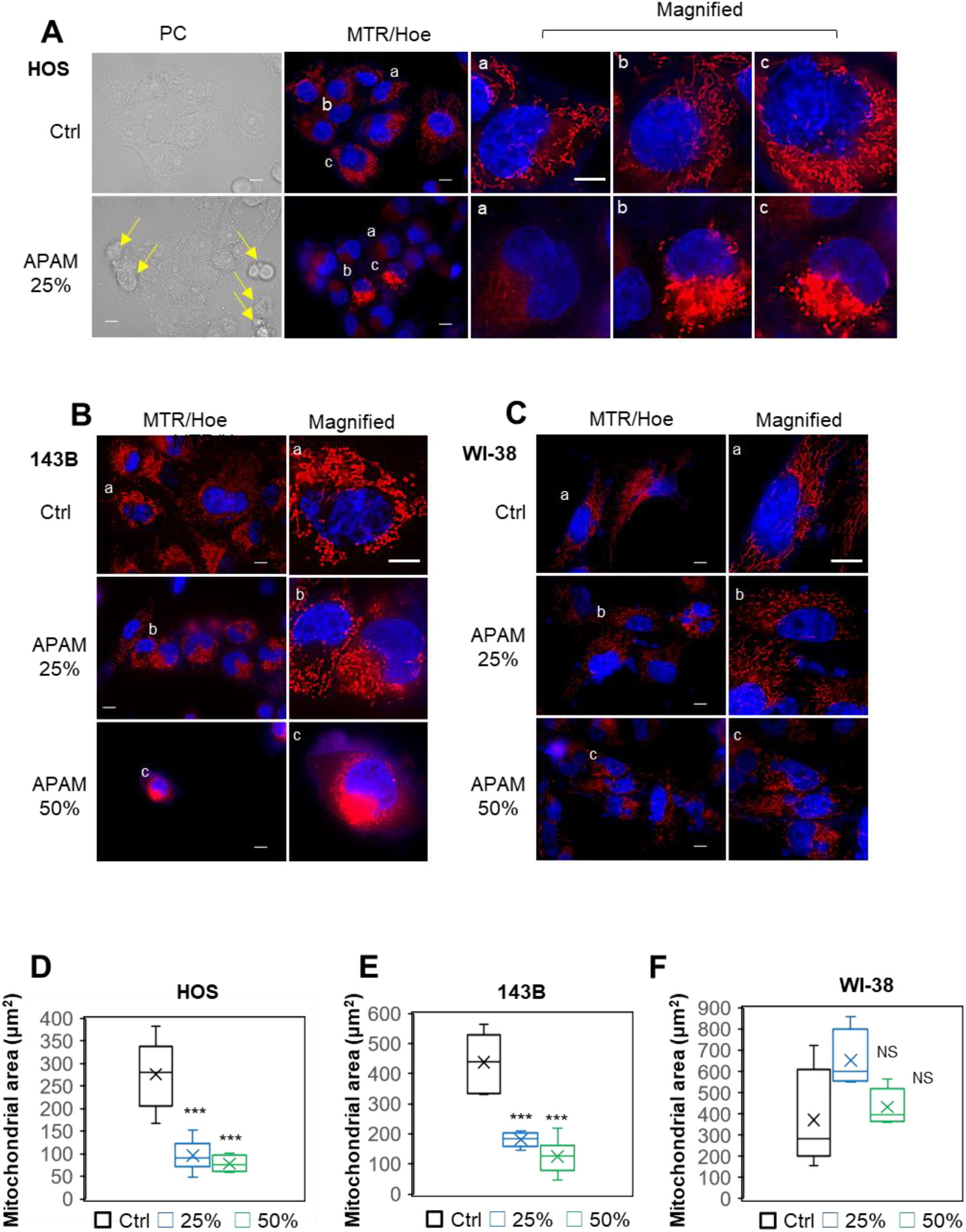
APAM induces MPMC in a tumor-selective manner. A‒C. After treating HOS (A), 143B (B), and WI-38 cells (C) with APAM (25, 50%) for 18 h, mitochondria and nuclei were stained with MTR and Hoechst 33342, respectively. Images were obtained and analyzed as described in the legend of Fig. 3. Magnified panels (a‒c) show the magnification of a‒c in the MTR/Hoe merged images. In A, the yellow arrows indicate damaged and dying cells. Bar = 10 μm. Note that in OS cells, MTR signals were dropped, fragmented, and clustered at the perinuclear sites after APAM treatment (A, B), while they became fragmented but minimally clustered in WI-38 cells (C). D‒F. HOS (D), 143B (E), and WI-38 cells (F) were treated with APAM (25, 50%) for 18 h, and mitochondria and nuclei were stained with MTR and Hoechst 33342 (Hoe), respectively. Images were taken as described in the legend of Fig. 3. Then, the occupied mitochondrial area in three different images per sample was measured using the BZ-H3A application as shown in Supplementary Fig. S4. Data are the mean ± SD (n = 4‒9). Data were analyzed by one-way analysis of variance followed by Tukey’s post hoc test. \*\*\**P <* 0.001; NS, not significant vs. control treated with vehicle.

### 2.6. MPMC occurs in a microtubule- and ROS-dependent manner

Mitochondria can move throughout the cytoplasm through the action of microtubules-associated motor proteins, Kinesin and Dynein (18, 32, 33). To determine the role of microtubules in MPMC, we attempted to analyze tubulin and mitochondrial dynamics simultaneously. After APAM treatment, the mitochondria and tubulin in HOS cells were stained with MTR and Oregon Green Paclitaxel, respectively, and observed microscopically. Like mitochondria, tubulin exhibited a network distributing broadly throughout the cytoplasm in untreated cells (Fig. 6A, top panels). Following APAM treatment, the tubulin network was mainly distributed to one side of the nuclei (Fig. 6A, second panels). Treatment with the microtubule inhibitor Nocodazole (NC) or the antioxidant NAC considerably abolished the tubulin network with minimal effects on tubulin’s subcellular distribution (Fig. 6A, third and fourth panels). Notably, NC and NAC blocked MPMC and tubulin redistribution (Fig. 6A, fifth and bottom panels). Similar results were obtained in 143B cells (Fig. 6B). These results indicate that MPMC occurs in a microtubule- and ROS-dependent manner.

**Fig. 6.**
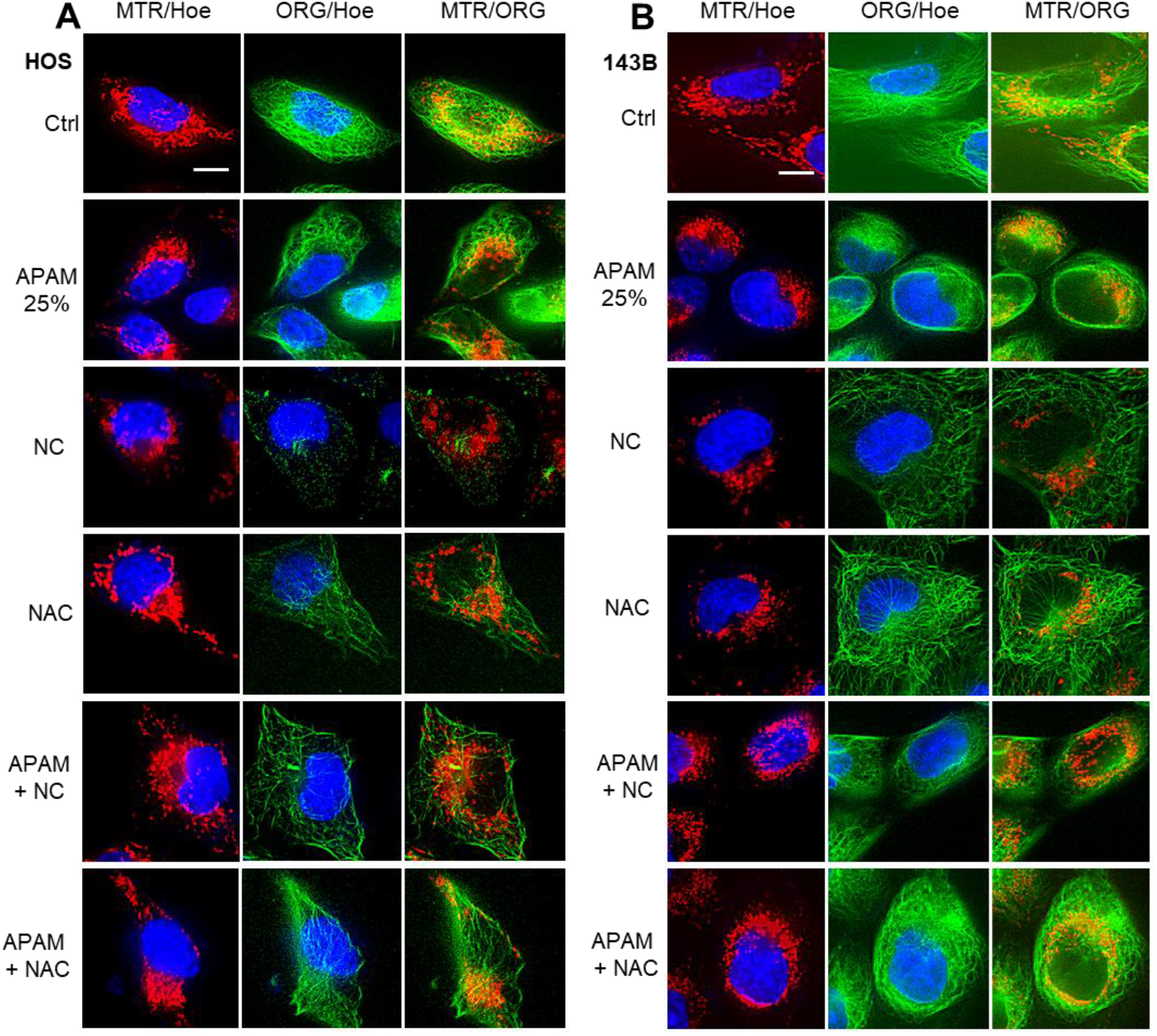
MPMC results from a microtubule- and ROS-dependent mitochondrial motility. HOS (A) and 143B cells (B) were pretreated with 100 nM NC or 2 mM NAC for 1 h and then treated with APAM (25%) for 2 h. Mitochondria, tubulin, and nuclei were stained with MTR, Oregon Green Paclitaxel (ORG), and Hoechst 33342 (Hoe), respectively. Images were taken as described in the legend of Fig. 3. Bar = 10 μm.

### 2.7. MPMC involves LPO accumulation and cardiolipin (CL) oxidation in the mitochondrial cluster

ROS can quickly attack lipids in the membranes, leading to lipid peroxides (LPOs) via peroxy radicals (ROO•). Therefore, we speculated that APAM might increase mitochondrial LPO (mLPO). To test this hypothesis, we analyzed the occurrence of LPOs using an LPO-reactive dye L248. Significant L248 puncta were distributed diffusely throughout the cytoplasm in untreated cells possessing healthy nuclei (Fig. 7A, upper panels). Notably, more concentrated L248 puncta were seen at the perinuclear sites in untreated cells containing smaller nuclei (indicated by yellow arrows). However, all L248 puncta were not colocalized with mitochondria in the cells. On the other hand, following APAM treatment, L248 puncta became a cluster at the perinuclear sites, colocalizing with mitochondria (Fig. 7A, lower panels). Since Cardiolipin (CL) is the major phospholipid found in the mitochondrial membrane and involved in cell death (34–36), APAM might increase its oxidation, leading to the production of the oxidized form of CL, CLOOH. The fluorescent dye 10-*N*-nonyl acridine orange (NAO) can bind to CL but not CLOOH (Petit et al., 1992) (37). Therefore, NAO staining enables us to judge the oxidative state of CL. We analyzed the mitochondrial NAO staining following APAM treatment. Substantial NAO signals entirely colocalizing with MTR signals were seen in untreated cells (Fig. 7B, top panels). APAM treatment resulted in marked concurrent decreases in MTR and NAO signals, indicating mitochondrial depolarization and CL oxidation (Fig. 7B, second panels). On the other hand, ferrostatin-1 (FS-1), a ROO• scavenger (38, 39), significantly increased NAO signals compared with untreated cells (Fig. 7B, third panels). Moreover, FS-1 prevented the decrease in NAO signals caused by APAM treatment (Fig. 7B, fourth panels). NAC treatment also inhibited the decrease in NAO signals while minimally affecting their intensity by itself (Fig. 7B, fifth and bottom panels). These results indicate that APAM can induce CL oxidation through oxidative stress via ROO•. Together, these data demonstrate that MPMC involves mLPO accumulation and CL oxidation.

**Fig. 7.**
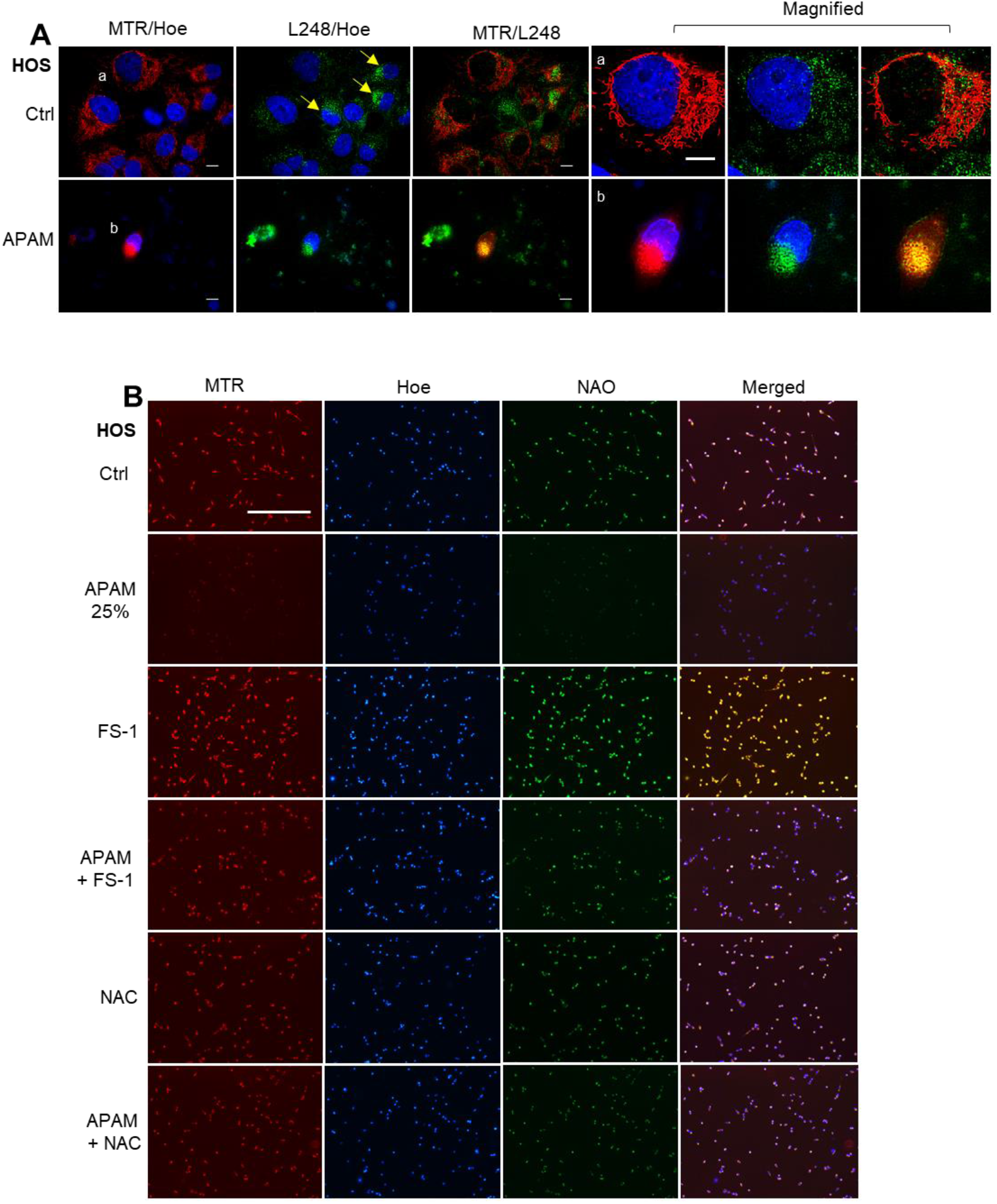
MPMC involves lipid peroxides (LPO) accumulation and Cardiolipin (CL) oxidation. A. HOS cells were treated with APAM (50%) for 2 h, and mitochondria, LPO, and the nuclei were stained with MTR, L248, and Hoechst 33342 (Hoe), respectively. Images were taken as described in the legend of Fig. 3. The magnified images (a, b) show the magnification of a, b in the merged images. Bar = 10 μm. B. HOS cells were pretreated with 1 μM FS-1 or 2 mM NAC for 1 h and then treated with APAM (25%) for 2 h, and mitochondria, CL, and the nuclei were stained with MTR, NAO, and Hoe, respectively. Images were taken and analyzed as described in the legend of Fig. 4. Bar = 400 μm.

### 2.8. APAM shows similar biological effects in OC cells

Next, we sought to determine whether these APAM’s effects were specific for OS or general in different cancers. Therefore, we examined whether APAM was also effective against OC because they belong to histologically distinct carcinoma from epithelial tissues. Results showed that APAM decreased the viability of SAS and HOC-313 cells in a dose-dependent manner (Fig. 8A, B). Moreover, Z-VAD-FMK significantly inhibited the effect of APAM (25%) but not APAM (50%) (Fig. 8C), indicating that the involvement of apoptosis and nonapoptotic cell death. APAM also induced MPMC in the cells in a dose-dependent manner (Fig. 8D). APAM also increased the mitochondrial O_2_^•-^, ^•^OH, and H_2_O_2_ in the cells, as shown by elevated MitoSOX, OxiOrange, and Hydrop signals, respectively (Fig. 8F). Additionally, live-cell imaging analysis using L248 revealed a substantial increase in LPO in the mitochondrial cluster in the cells (Fig. 8G). Together, these results indicate that APAM induces cell death and MPMC in OC through possibly similar mechanisms.

**Fig. 8.**
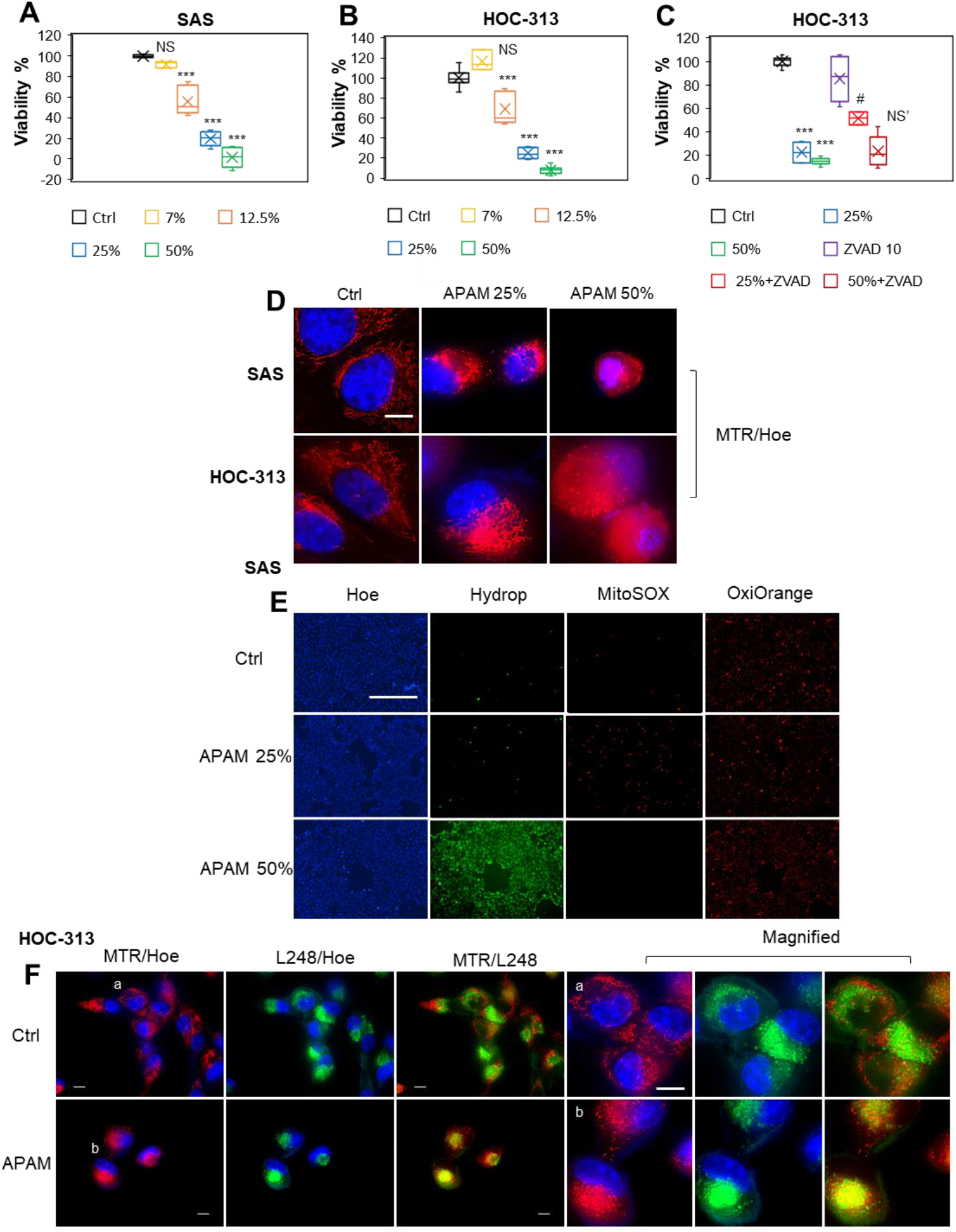
APAM induces cell death, ROS generation, LPO accumulation, and MPMC in OC. A‒C. SAS (A), HOC-313 (B) cells (4 × 10^3^ cells) were treated with the indicated concentrations of APAM for 72 h and analyzed for viability using a WST-8 cell growth assay. C. HOC-313 cells were pretreated with 10 μM Z-VAD-FMK for 1 h and then treated with APAM (25, 50%). Data are the mean ± SD (n = 7). Data were analyzed by one-way analysis of variance followed by Tukey’s post hoc test. \*\*\**P <* 0.001; NS, not significant vs. control treated with vehicle. #*P <* 0.05; NS’, not significant vs. APAM alone. D. After treating SAS (upper) and HOC-313 cells (lower) with APAM (25, 50%) for 18 h, mitochondria and nuclei were stained with MTR and Hoechst 33342 (Hoe), respectively. MTR/Hoe merged images were obtained and analyzed as described in the legend of Fig. 3. Bar = 10 μm. E. SAS cells (1.5×10^4^) were treated with APAM (25, 50%) for 2 h and then stained with 5 μM MitoSOX, 1 μM Hydrop, or 1 μM OxiOrange. The nuclei were counterstained with Hoe. Images were obtained and analyzed as described in the legend of Fig. 4. Bar = 400 μm. F. HOC-313 cells were treated with APAM (50%) for 2 h, and mitochondria, LPO, and the nuclei were stained with MTR, L248, and Hoe, respectively. Images were taken as described in the legend of Fig. 3. The magnified images (a, b) show the magnification of a, b in the merged images. Bar = 10 μm.

### 2.9. APAM induces minimal cell death, mROS generation, mLPO accumulation, CL oxidation, and MPMC in fibroblasts

Given that MPMC is a prodeath event, one can suppose that it could not occur in normal cells where APAM has no cytotoxic activity. To test if this was the case, we examined the effect of APAM on the mitochondrial network and positioning in fibroblasts. Minimal cell and nuclear damages occurred following APAM treatment for 24 h (Fig. 9A). APAM substantially increased mitochondrial fragmentation but minimally evoked MPMC (Fig. 9B). APAM induced minimal increases in O_2_^•-^, LPO, and CL oxidation in the mitochondria (Fig. 9A, C, D). These results indicate that mROS generation, mLPO accumulation, CL oxidation, and MPMC occur in parallel cell death in a tumor-selective manner.

**Fig. 9.**
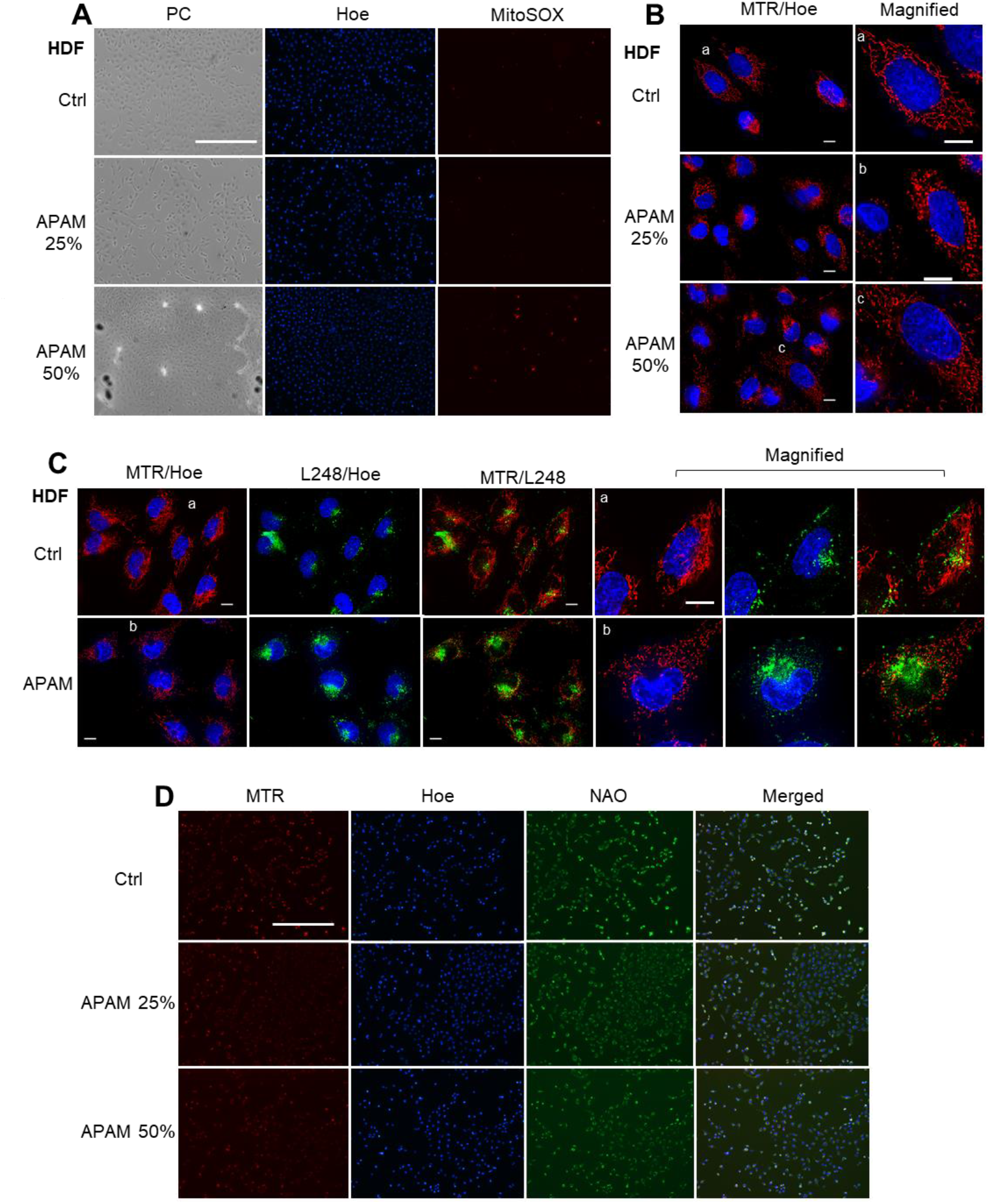
APAM causes minimal cell death, ROS generation, LPO accumulation, CL oxidation, and MPMC in normal cells. A. HDF cells (1.5×10^4^) were treated with APAM (25, 50%) for 2 h and then stained with 5 μM MitoSOX and Hoechst 33342 (Hoe). Images were obtained and analyzed as described in the legend of Fig. 4. PC, phase contrast. Bar = 400 μm. Note that minimal cell morphological changes were observed in APAM-treated cells. B. After treating the cells with APAM (25, 50%) for 18 h, mitochondria and nuclei were stained with MTR and Hoe, respectively. Images were obtained and analyzed as described in the legend of Fig. 3. Bar = 10 μm. Note that minimal mitochondrial clustering and nuclear damages occurred in APAM-treated cells. C. The cells were treated with APAM (50%) for 2 h, and mitochondria, LPO, and the nuclei were stained with MTR, L248, and Hoe, respectively. Images were taken as described in the legend of Fig. 3. The magnified images (a, b) show the magnification of a, b in the merged images. Bar = 10 μm. D. The cells were treated with APAM (25, 50%) for 2 h, and mitochondria, CL, and the nuclei were stained with MTR, NAO, and Hoe, respectively. Images were taken and analyzed as described in the legend of Fig. 4. Bar = 400 μm.

## 3. Discussion

In the present study, we investigated the antitumor activity of APAM and its possible impact on the mitochondrial network and subcellular positioning. APAM had potent antitumor activity against OS (Fig. 1) and OCs (Fig. 8) *in vitro.* In contrast, APAM had minimal cytotoxicity against human dermal and lung fibroblasts, indicating that it acts in a tumor-selective manner. In supporting these results obtained *in vitro,* APAM significantly reduced allograft or xenograft OS tumors with minimal adverse effects (Fig. 1). These observations indicate that APAM has a tumor-selective cytotoxic activity, like other various PAM prepared with different gasses or culture media (11‒16). It is essential to know the mechanisms underlying antitumor action. APAM could activate apoptosis and nonapoptotic cell death depending on the cell line (Fig. 2). Notably, the antitumor activity of APAM (25%) was substantially inhibited by Z-VAD-FMK, while that of APAM (50%) was minimally affected by it (Fig. 2 and 8). These observations indicate that APAM can switch cell death modality from apoptosis to nonapoptotic cell death depending on the concentration. These properties are similar to those of H_2_O_2_ (40) and oxidative phosphorylation (OXPHOS) inhibitors (41). Both H_2_O_2_ and OXPHOS inhibitors such as FCCP and antimycin A can activate apoptosis and nonapoptotic cell death depending on the concentration and processing time. Specifically, cell death during the early 24 h was sensitive to Z-VAD-FMK, while that observed during another 48 h was insensitive (41). Highly proliferating cancer cells could rapidly consume nutrients in medium, resulting in an energy-deficient status during prolonged processing. Accordingly, they could be more prone to necrotic cell death than apoptosis. Like APAM, H_2_O_2_ and OXPHOS inhibitors can evoke mitochondrial O_2_^•-^generation and have antitumor activities through oxidative stress (40, 41). Together, mitochondrial oxidative stress may lead to different cell death pathways depending on the intensity and duration. Moderate and short stress can activate apoptosis, while intense and persistent stress may be required for nonapoptotic cell death in apoptosis-resistant cells.

Our results indicate that APAM can activate substantial autophagic flux (Fig. 3). The observations are similar to the previous reports that demonstrate the role of autophagic cell death in PAM’s effects (14, 42). Autophagy copes with energy demands, while mitophagy is essential for removing damaged organelles. Accordingly, autophagy may be particularly critical for the survival of cancers in which high energy demands and constant removal of damaged mitochondria are required. On the other hand, intense and persistent autophagy activation results in autophagic cell death, which might act as a tumor-suppressive event (43, 44). Notably, like HePAM (14), APAM also induced mitophagy in tumor cells but not fibroblasts (Fig. 3), indicating the role in APAM’s cytotoxic activity. Although the role of mitophagy remains unclear, it might play a role in mitochondrial fragmentation because APAM induced mitochondrial depolarization, a potent cause of mitophagy (22). Ferroptosis, the iron-mediated necrotic cell death, has emerged as another cell death modality activated in various cancers by anticancer compounds (38, 39). However, given that no mitochondrial requirement for ferroptosis (45), APAM-induced cell death seems different from ferroptosis. The roles of autophagy and ferroptosis in APAM-induced cell death are currently under investigation in our laboratory.

It has been long known that mitochondria are highly plastic, dynamic, and heterogenous organelles. Their size and shape (network shape) vary in different tissues, cells, and experimental conditions. However, only recently has much attention been paid to the biological significance of their heterogeneous properties. An emerging view is that changes in mitochondrial network and location are not merely passive events but active events coupling with cellular functions and survival. Concerted mitochondrial dynamics and subcellular positioning are essential for cell function, survival, and specific cancer onset and progression (24, 25). Mitochondria can distribute throughout the cytoplasm (pan-cytoplasmic), subplasmalemmal, or perinuclear. Precise distributions are essential for energy supply (18, 19), the plasma membrane Ca^2+^ channel activity (46), Ca^2+^ signaling (47‒49), and hypoxic state and signaling (26‒29). Moreover, several reports demonstrate a closed relationship between mitochondrial motility and the mitochondria-ER tethering (50), Ca^2+^ transport (51), and redox state (52, 53). Our previous works show that mitochondrial fragmentation followed by clustering is essential for apoptotic or nonapoptotic cell death caused by TRAIL, HePAM, and PAI (12, 13, 17, 30, 31). All these substances modulate the mitochondrial network in tumor cells specifically. HePAM activates both Drp1-dependent and -independent mitochondrial fragmentation, and exogenously added H_2_O_2_ mimics the former mechanism (12). The present study revealed that APAM also evoked mitochondrial fragmentation and clustering in a tumor-selective manner (Fig. 5). APAM and HePAM (12, 13) are chemically and biologically similar. They commonly have H_2_O_2_, increasing mitochondrial O_2_^•-^, inducing mitochondrial depolarization and activating autophagy and mitophagy. Notably, in normal cells, both HePAM and H_2_O_2_ (12, 13) and APAM (Fig. 3 and 9) minimally increased mitochondrial O_2_^•-^ and mitophagy. Additionally, HePAM and H_2_O_2_ can activate Drp1 phosphorylation at Ser 616, the driving force to mitochondrial fission in tumor cells but not normal cells (12, 13). Despite the facts, mitochondrial fragmentation caused by HePAM is augmented but not suppressed by the Drp1 inhibitor Mdivi-1 or Drp1 gene silencing (12). Collectively, an intriguing possibility is that mitochondrial fragmentation is triggered by the Drp1-dependent fission pathway via mitochondrial oxidative stress and exacerbated by another Drp1-independent pathway where autophagy and mitophagy might play an additional role. Further examination of this scenario is underway.

Moreover, we discovered that the mitochondrial network collapse eventually led to a unique mitochondrial subcellular positioning, MPMC. This event occurs in all tumor cell lines tested (Fig. 5 and 8) but not lung and dermal fibroblasts (Fig. 5 and 9). MPMC occurred within hours after APAM treatment and was prevented by NAC (Fig. 6). Thus, it may be a prodeath event where ROS play a role. At present, the precise mechanisms underlying MPMC remain unclear. However, the simultaneous occurrence of mROS (Fig. 4 and 8), mLPO accumulation, and CL oxidation and prevention by FS-1 (Fig. 7 and 8) indicate the critical role of mLPO accumulation. Moreover, concomitant translocation of the mitochondrial and tubulin networks and their prevention by NC (Fig. 6) strongly suggest the involvement of mitochondrial motility through the action of microtubules-associated motor proteins. Together, our data coincide with the working model shown in Fig. 10. APAM increases mROS, including O_2_^•-^, H_2_O_2_, and ^•^OH, promoting nonphysiological mitochondrial fragmentation, possibly via intensive and persistent mitophagy. Then, the fragmented mitochondria will move along with microtubules and gather at one side of the perinuclear sites. This working model raises several interesting questions. For instance, how MPMC leads to cell death, why it occurs tumor-selectively, and why mitochondria gather unipolar perinuclear regions. Since MPMC is pronounced explicitly in cells possessing shrank and decomposed nuclei, it might play a role in nuclear damages. Given the critical role of PNMC in hypoxia, MPMC also might compromise hypoxic status in cancer cells. PNMC has been shown to contribute to the stabilization of HIF-1α. Mitochondria in PNMC can generate ROS (mROS), which in turn increases nuclear ROS (nROS), leading to oxidation-mediated HIF-1α stabilization and the downstream hypoxia signaling (26–29). In addition, the activated metabolism and genetic instability under the control of oncogenic transformations cause increased ROS generation and decreased antioxidant systems in cancer cells. Thus, cancer cells may be exposed to higher oxidative stress than normal cells. Therefore, APAM may preferentially make tumor cells exceed the allowed mitochondrial oxidative stress range and undergo MPMC. Finally, we are currently attempting to know the biochemical characteristics of the perinuclear sites in which mitochondria make clusters. In summary, the present study shows that the tumor-targeting agent, APAMcan switch the mitochondrial positioning from PNMC to MPMC in OS and OC cells but not fibroblasts. Cancer cells seem to be more prone than normal cells to the switch due to their intense ambient oxidative stress. Thus, MPMC might serve as a new promising objective for tumor-targeting anticancer compounds.

**Fig. 10.**
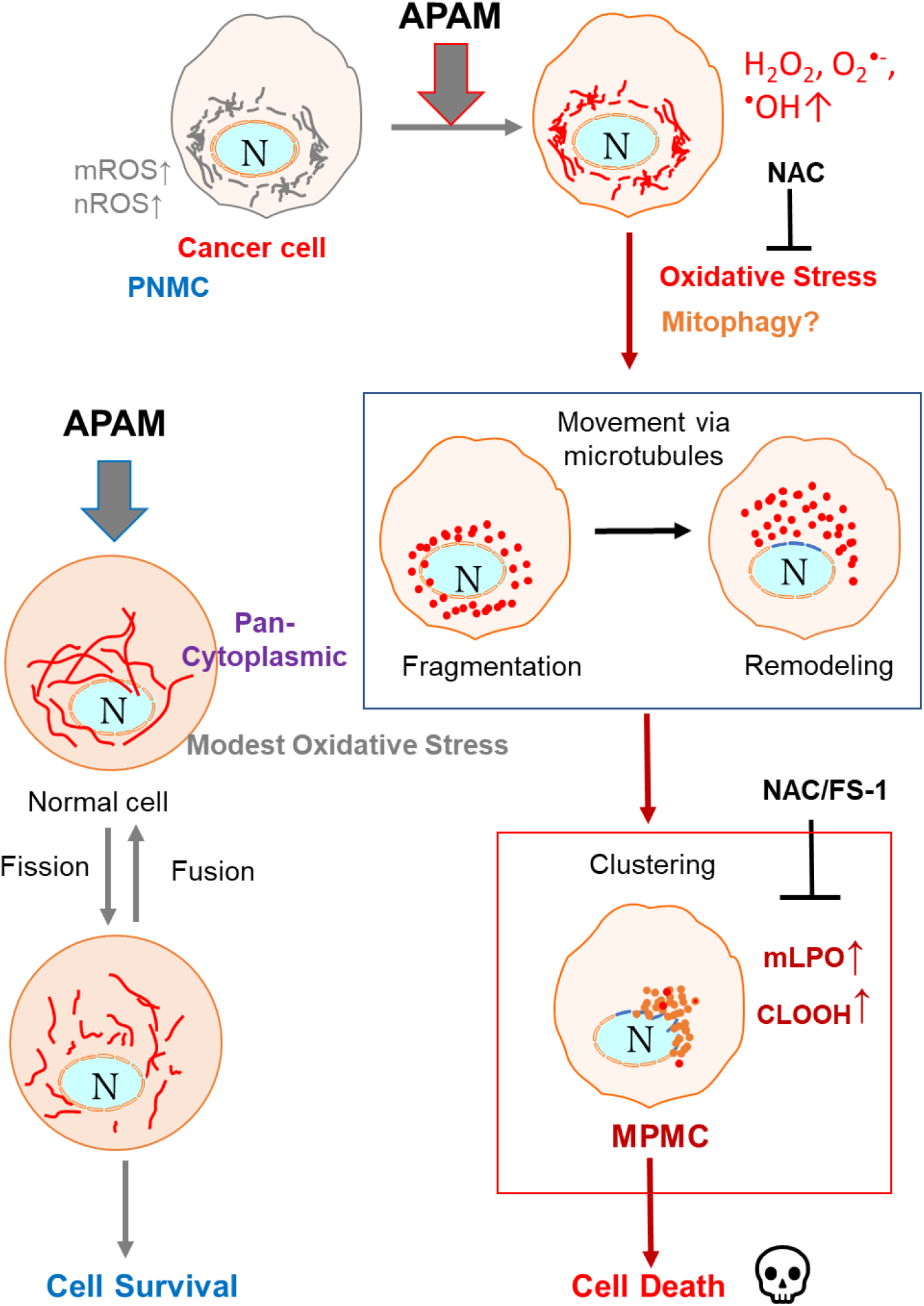
The working model in this study. Cancer cells are adapting to hypoxic microenvironments owing to PNMC. Mitochondria in PNMC can generate ROS (mROS), which increases nuclear ROS (nROS), oxidation-mediated HIF-1α stabilization, and the downstream hypoxia signaling. APAM induces massive mROS and a nonphysiological mitochondrial fragmentation mechanism, possibly intensive and persistent mitophagy. The fragmented mitochondria will move along with microtubules and form clusters at one side of the perinuclear sites (MPMC), leading to cell death. On the other hand, normal cells possess pan-cytoplasmic mitochondria, and APAM increases mitochondrial oxidative stress only modestly. It causes the physiological Drp1-dependent mitochondrial fission, reversible through the fission-fusion dynamics, leading to cell survival.

## 4. Materials and methods

### 4.1. Materials

All chemicals were purchased from Sigma Aldrich (St. Louis, MO, USA) unless otherwise specified. The pan-caspase inhibitor Z-VAD-FMK was purchased from Merck Millipore (Darmstadt, Germany). All insoluble reagents were dissolved in dimethyl sulfoxide (DMSO) and diluted with high glucose-containing Dulbecco’s modified Eagle’s medium (DMEM) supplemented with 10% fetal bovine serum (FBS) or Hank’s balanced salt solution (HBSS; pH 7.4, Nissui Pharmaceutical Co., Ltd., Tokyo, Japan) (final DMSO concentration, < 0.1%) before use.

### 4.2. Cell Culture

The human OC cell line, SAS (JCRB0260) was obtained from the Japanese Collection of Research Bioresources (JCRB) Cell Bank of National Institutes of Biomedical Innovation, Health, and Nutrition (Osaka, Japan). Another human OC cell line, HOC313, was kindly provided by the Department of Oral and Maxillofacial Surgery, Graduate School of Medical Science, Kanazawa University (Kanazawa, Japan). Human fetal lung fibroblasts, WI-38 (JCRB9017), were obtained from JCRB. Human dermal fibroblasts (HDFs) from the facial dermis were obtained from Cell Applications (San Diego, CA). Human osteosarcoma HOS (RCB0992), 143B (RCB0701), and murine osteosarcoma LM8 (RCB1450) cells were purchased from Riken Cell Bank (Tsukuba, Japan). The cells were maintained in 10% FBS (GIBCO^®^, Life Technologies) containing DMEM (GIBCO^®^, Life Technologies, Carlsbad, CA, USA) (FBS/DMEM) supplemented with 100 U/mL penicillin and 100 μg/mL streptomycin at 37°C in a humidified atmosphere with 5% CO_2_.

### 4.3. APAM preparation

CAP was generated from the ambient air using a Piezobrush*™* PZ2 model plasma jet (relyon, Germany) equipped with a piezo element. The typical experimental conditions are a frequency of >50 kHz, a voltage of >20 kV, and an electron density of 10^14∼16^/cm. APAM (1 mL) was made by irradiating plasma from above at a distance of 20 mm to 1 mL of DMEM without phenol red for 1 min. The original APAM was diluted to a final concentration of 6.3–50% with 10% FBS/DMEM (for cell experiments) or HBSS (for biochemical experiments) and was indicated as APAM (6.3–50%).

### 4.4. Cell viability assay

Cell viability was measured by the WST-8 assay using Cell Counting Reagent SF (Nacalai Tesque, Inc., Kyoto, Japan) or Cell Counting Kit-8 (Dojindo Molecular Technologies, Inc., Kumamoto, Japan) according to the manufacturer’s instructions. These methods are colorimetric assays based on the formation of a water-soluble formazan product (WST-8). Briefly, cells were seeded at a density of 3 or 4×10^3^ cells/well in 96-well plates (Corning Incorporated, Corning, NY, USA). They were cultured with agents to be tested for 72 h at 37°C before adding 10 μL cell counting reagent and further incubation for 2 h. Absorbances at 450 nm were measured using a Nivo (PerkinElmer Japan Co., Ltd., Yokohama, Japan) or SH-1100R (Lab) microplate reader (Corona Electric Co., Ltd., Ibaraki, Japan).

### 4.5. Animals and *in vivo* antitumor activity

All animal experiments were conducted according to the Guidelines for Proper Conduct of Animal Experiments, Science Council of Japan. Protocols were approved by the Animal Care and Use Committee (No.17–11), University of Yamanashi. Male C3H/HeJJcl mice and BALB/cAJcl-nu/nu mice were purchased from CLEA Japan, Inc. (Tokyo, Japan). The mice were housed at 22–24°C under a 12-h light/dark cycle and were fed standard mouse chow. Water was available ad libitum. The ability of APAM to reduce tumor growth *in vivo* was evaluated using allograft transplants of LM8 cells in mice. Male C3H/HeJJcl or BALB/cAJcl-nu/nu mice (8 weeks of age) were administered general anesthesia with isoflurane (ISOFLU; Abbott Laboratories, North Chicago, IL, USA) and oxygen. LM8 cells (1×10^6^ cells/mouse) in 100 µL DMEM were injected subcutaneously into the backs of the CH3 mice on day 0. Alternatively, 143B cells (1×10^6^ cells/mouse) in 100 µL of DMEM were injected intramedullary into the right tibia of the BALB/cAJcl-nu/nu mice on day 0. On day seven post-transplant, 200 μL of APAM or vehicle were administered intravenously to six mice in each group every other day. The mice were weighed, and the primary tumors were measured weekly. A schematic diagram of the experiment is shown in Supplementary Fig. S1.

### 4.6. Cell death assay

Cells were cultured in 6-well plates for 24 h and then exposed to APAM (50%) for 24 h. The cells were retrieved using Versene (GIBCO®, Life Technologies) and incubated with APC-conjugated Annexin V and 7-AAD (BD Biosciences) for 15 min to evaluate apoptotic cell death. Data were collected using a FACSCelesta™ flow cytometer (BD Biosciences, Franklin Lakes, NJ, USA). Data obtained were analyzed by CellQuest Pro (Becton Dickinson Biosciences) and FlowJo software (TreeStar, Ashland, OR). Experiments were performed in triplicate. Annexin V-positive cells were defined as apoptotic cells.

### 4.7. Western blotting analysis

LM8 and 143B cells were collected, and cell lysates were prepared using a CelLytic MT cell lysis reagent (Sigma-Aldrich) according to the manufacturer’s instructions. Western blotting analysis was performed as previously reported (17). Briefly, equal amounts of protein from each sample were analyzed by immunoblotting with primary antibodies against cleaved caspase-3 (Asp175) (5A1E) (#9664, 1:1000), caspase-3 (#9662, 1:1000), LC3-A/B (D3U4C) (#12741, 1:1000), LC3-B (D11) (#3368, 1:1000), and GAPDH (D16H11) (#5174, 1:1000) (Cell Signaling Technology, Danvers, MA, USA). Images were captured using a LAS-4000 camera system from Fujifilm (Tokyo, Japan) and quantified using ImageJ 1.52a (Wayne Rasband, National Institutes of Health, Bethesda, MD, USA).

### 4.8. Mitochondrial network and positioning assays

The mitochondrial network in live cells was analyzed as previously described (17). Briefly, cells in FBS/DMEM (3×10^4^/well) adherent on 8-well chambered coverslips were treated with the agents to be tested for 24 h at 37°C *at a* 5% CO_2_ incubator. After removing the medium by aspiration, the cells were washed with fresh FBS/DMEM and stained with 20 nM MitoTracker^TM^ Red CMXRos (MTR) for 1 h at 37°C at *the* CO_2_ incubator. In positioning experiments, the nuclei were counterstained with 1 mg/mL of Hoechst 33342. The cells were then washed and immersed in FluoroBrite^TM^ DMEM (Thermo Fisher Scientific, Waltham, MA., USA). Images were obtained using a BZ X-7*1*0 Digital Biological Microscope (Keyence, Osaka, Japan) equipped with a 100 ×, 1.40 n.a. UPlanSApo Super-Apochromat, coverslip-corrected oil objective (Olympus, Tokyo, Japan) and analyzed using BZ-H3A application software (Keyence). The occupied mitochondrial area was measured in three photos per sample using the BZ-H3M application, as shown in Supplementary Fig. S3.

### 4.9. Autophagy assay

Cells were cultured in 6-well plates for 18 h and then exposed to APAM (50%) for 24 h. Cells were stained and analyzed using DALGreen (Dojindo, Kumamoto, Japan) according to the manufacturer’s instructions. Data were collected using a FACSCelesta™ flow cytometer (BD Biosciences, Franklin Lakes, NJ, USA) and analyzed using FlowJo software (TreeStar Inc., Palo Alto, CA, USA). Experiments were performed in triplicate. In mitophagy experiments, mitochondria and autophagosomes were stained with MTR and the specific dye Cyto-ID® (Enzo Life Sciences), respectively. Mitophagy was judged from colocalization between mitochondria and the Cyto-ID signals.

### 4.10. Quantitation of H_2_O_2_ in APAM

The concentration of H_2_O_2_ in APAM was measured using the Amplex Red Hydrogen Peroxide/Peroxidase Assay Kit (Thermo Scientific, Rockford, IL, USA) according to the manufacturer’s protocols as previously described (17). Briefly, samples were diluted appropriately and placed on a 96-well plate (50 μL/well). Then, 50 μL of a working solution of 100 μM of Amplex Red reagent and 0.2 U/mL of horseradish peroxidase was added to the well and incubated at room temperature for 30 min. The absorbance at 570 nm was measured using a Nivo 3F Multimode Plate Reader (PerkinElmer Japan Co., Ltd., Yokohama, Japan). The concentrations of H_2_O_2_ were calculated using a standard curve made using the authentic H_2_O_2_ in the Kit.

### 4.11. Measurements of intracellular ROS generation

Cells (1.5×10^4^/well) in FBS/DMEM were cultured on a 35-mm poly-Lysine-coated glass-bottom dish (Matsunami Glass, Tokyo, Japan), treated with the agents to be tested for 2 h at 37°C at a CO_2_ incubator. After removing the medium by aspiration, the cells were placed in FluoBrite^TM^ DMEM and stained with 5 μM MitoSOX^TM^ Red (MitoSOX, Thermo Fisher Scientific), 1 μM OxiOrange^TM^ or 1 μM Hydrop^TM^ (Goryo Chemicals, Sapporo, Japan) for 20 min. Images were obtained with EVOS FL Cell Imaging System (Thermo Fisher Scientific) equipped with a 10 ×objective and analyzed using the freely available NIH ImageJ software (NIH, Bethesda, MD, USA) as previously described (30).

### 4.12. LPO and CL oxidation analyses

Cells (3×10^4^/well) were cultured and treated as described above and stained with 20 nM MitoTracker Red CMXRos, 1 μM L248 (Liperfluo, Dojindo), or 100 nM 10-*N*-Nonyl acridine orange (NAO, Sigma-Aldrich) for 20 min. Images were obtained with BZ X-710 Digital Biological Microscope (Keyence) or EVOS FL Cell Imaging System (Thermo Fisher Scientific). They were analyzed using BZ-H3A/H3M application (Keyence) or NIH ImageJ software (NIH) as described above. LPO accumulation in mitochondria was judged by colocalization between MTR and L248 signals in merged images.

### 4.13. Statistical analysis

Data are presented as mean ± standard deviation (SD) and were analyzed by one-way analysis of variance followed by Tukey’s post hoc test using an add-in software with Excel 2016 for Windows (SSRI, Tokyo, Japan). For some experiments, significance was determined using Student’s t-test after an F-test. P < 0.05 was considered statistically significant.

## Acknowledgments

We thank the JCRB Cell Bank of National Institutes of Biomedical Innovation, Health, and Nutrition (Osaka, Japan) and the Riken BioResource Center (Tsukuba, Japan) for providing cell lines. We thank M Ubagai and C Chino for their technical assistance.

## Supplementary Figure Legends

**Supplementary Fig. S1.**
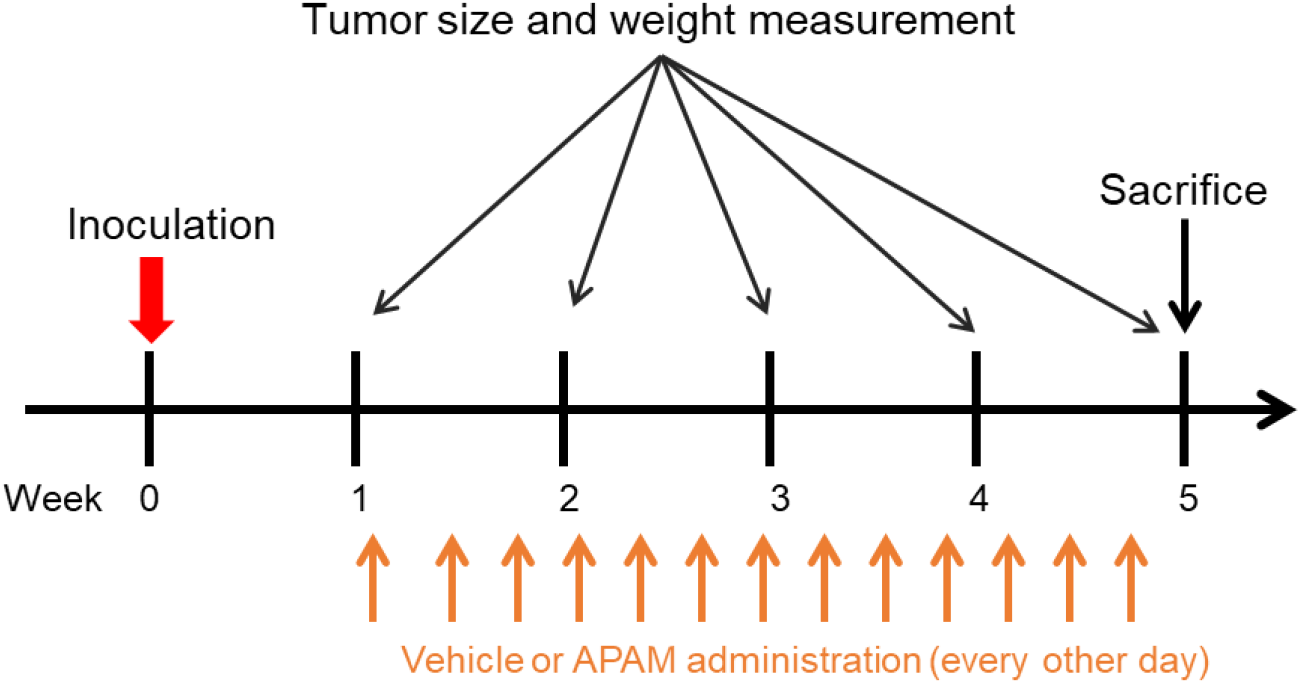
Schematic diagram showing the experimental design for *in vivo* antitumor activity assay. Cells (1 × 10^6^ cells/mouse) in 0.1 ml DMEM were inoculated on day 0. From day 7, APAM (50%) was administered three times per week intravenously to 6 mice per group. The mice were weighed, and the size of the primary tumors was measured every week. On day 35, all mice were sacrificed.

**Supplementary Fig. S2.**
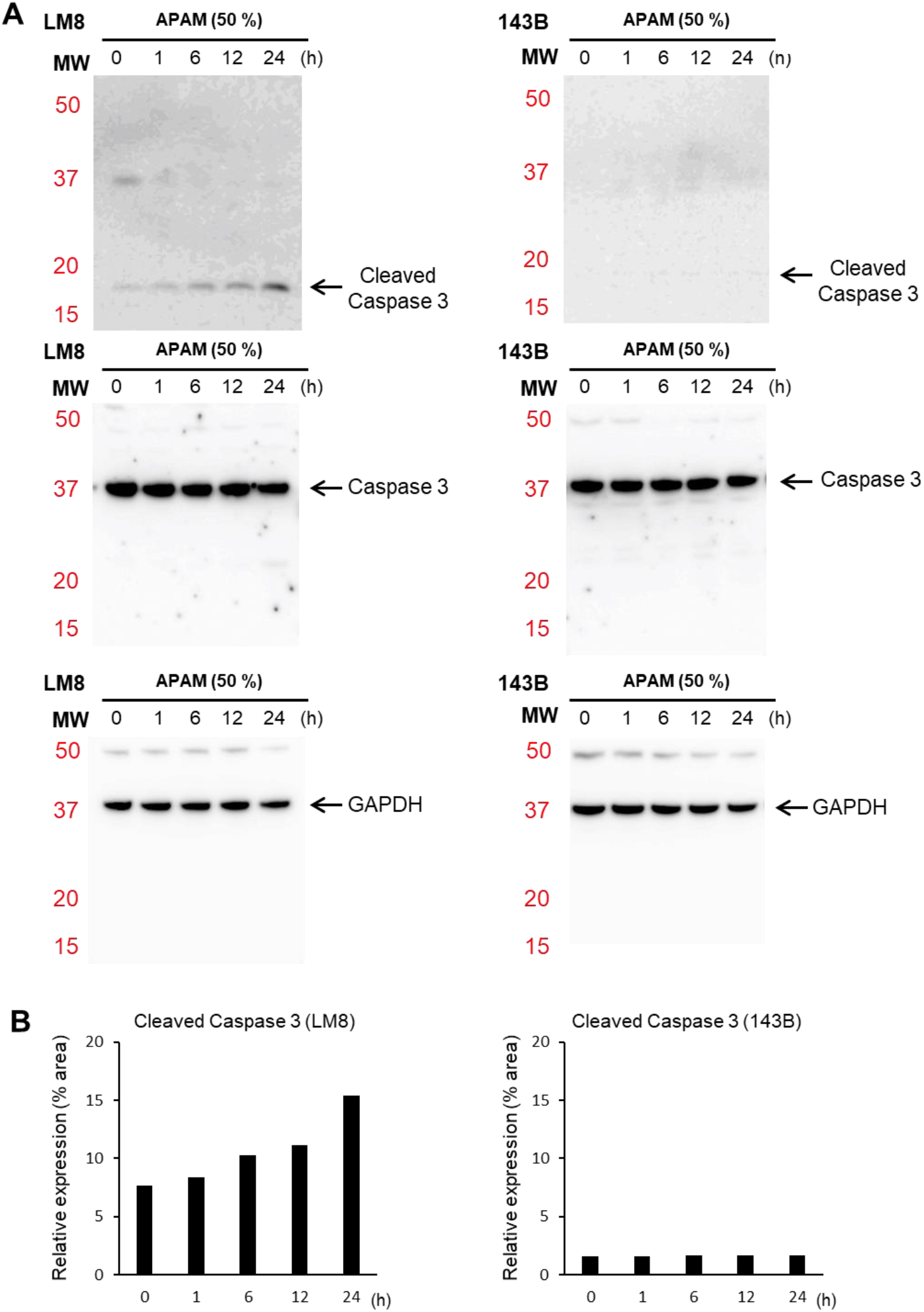
A, B. Examples of uncropped western blots (A) and quantification (B) for cleaved caspase-3 and GAPDH in Fig. 2C, D.

**Supplementary Fig. S3.**
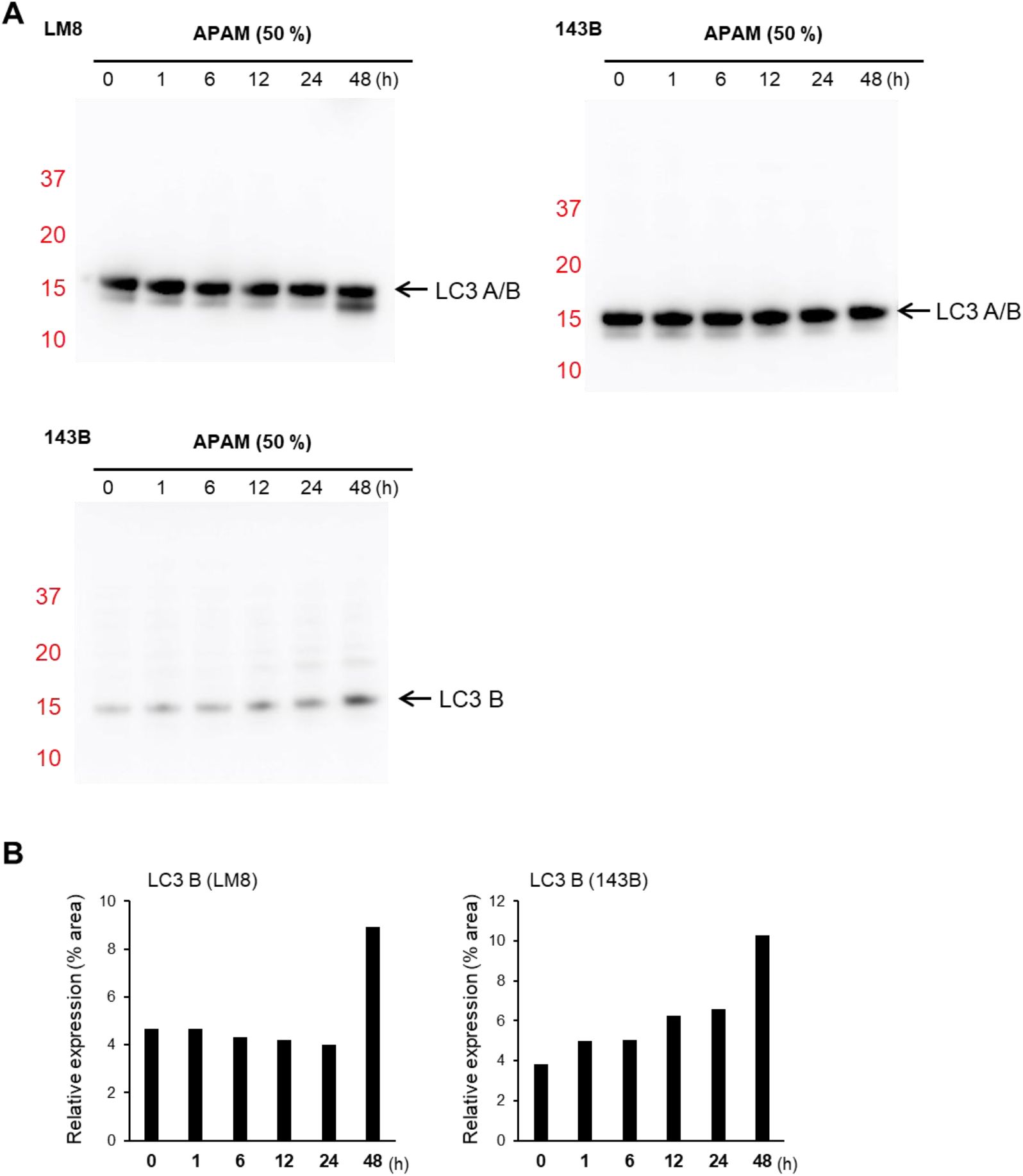
A, B. Examples of uncropped western blots (A) and quantification (B) for LC3-A/B, LC-3B, and GAPDH in Fig. 3C.

**Supplementary Fig. S4.**
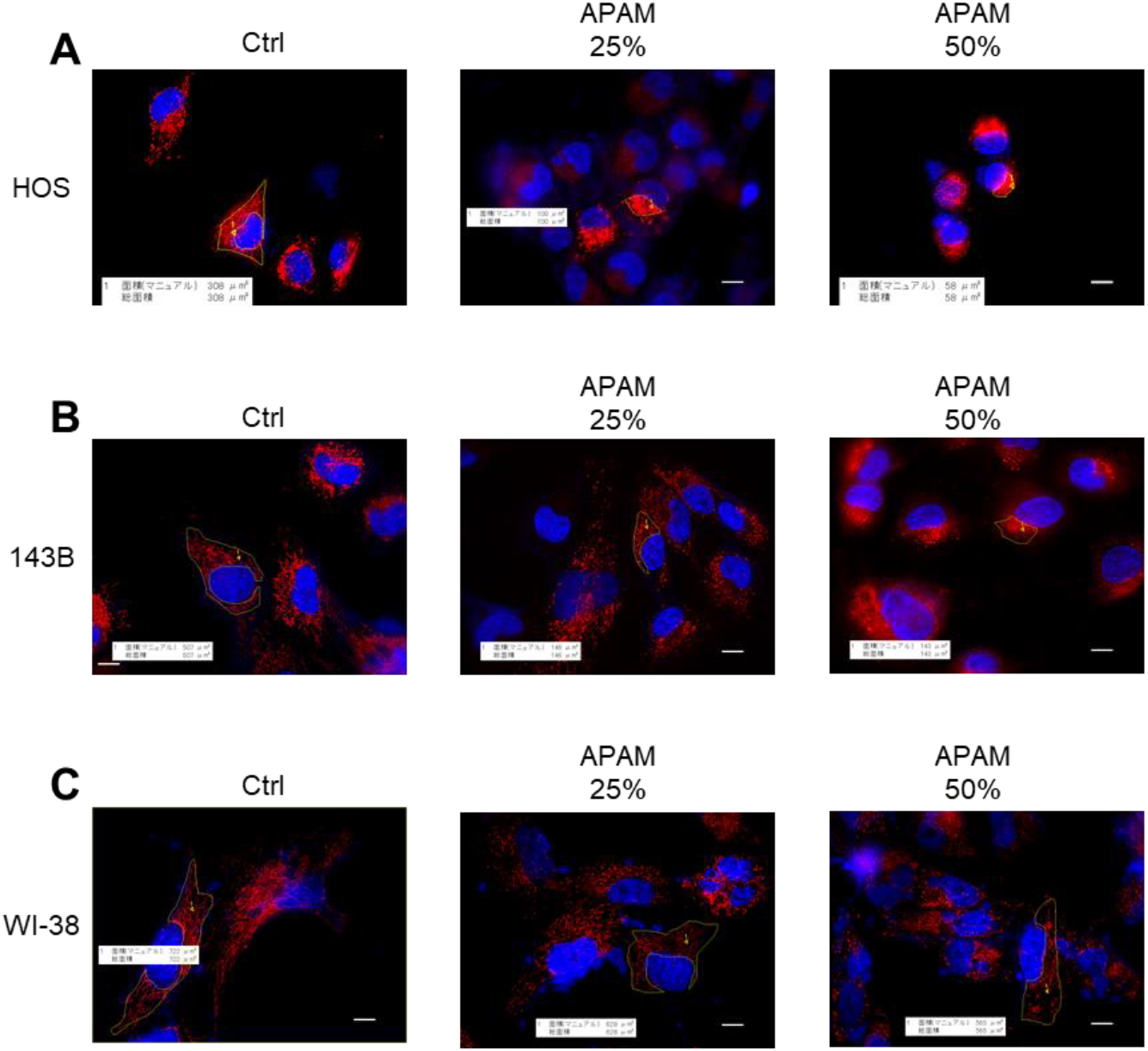
Examples of the measurements of the occupied mitochondrial area (OMA). HOS (A), 143B (B), and WI-38 cells (C) were treated with APAM (25, 50%) for 18 h, and mitochondria and the nuclei were stained with MitoTracker Red CMXRos (red) and Hoechst 33342 (blue), respectively. Images were obtained using a BZ710 Digital Biological Microscope equipped with a 100 × coverslip-corrected oil objective and analyzed using the BZ-H3A application. OMA was manually selected in three pictures indicated by the yellow line and automatically measured using the BZ-H3M application. Bar = 10 μm.

## References

1. Luetke A, Meyers PA, Lewis I, Juergens H. Osteosarcoma treatment - where do we stand? A state of the art review. Cancer Treat Rev. 2014;40(4):523–32. doi: 10.1016/j.ctrv.2013.11.006.

2. Siegel RL, Fedewa SA, Miller KD, Goding-Sauer A, Pinheiro PS, Martinez-Tyson D, et al. Cancer statistics for Hispanics/Latinos, 2015. CA Cancer J Clin. 2015;65(6):457–80. doi: 10.3322/caac.21314.

3. Morrow JJ, Bayles I, Funnell APW, Miller TE, Saiakhova A, Lizardo MM, et al. Positively selected enhancer elements endow osteosarcoma cells with metastatic competence. Nat Med. 2018;24(2):176–85. doi: 10.1038/nm.4475.

4. Glick D, Barth S, Macleod KF. Autophagy: cellular and molecular mechanisms. J Pathol. 2010;221(1):3–12. doi: 10.1002/path.2697.

5. Kornienko A, Mathieu V, Rastogi SK, Lefranc F, Kiss R. Therapeutic agents triggering nonapoptotic cancer cell death. J Med Chem. 2013;56(12):4823–39. doi: 10.1021/jm400136m.

6. Keidar M, Walk R, Shashurin A, Srinivasan P, Sandler A, Dasgupta S, et al. Cold plasma selectivity and the possibility of a paradigm shift in cancer therapy. Br J Cancer. 2011;105(9):1295–301. doi: 10.1038/bjc.2011.386.

7. Zucker SN, Zirnheld J, Bagati A, DiSanto TM, Des Soye B, Wawrzyniak JA, et al. Preferential induction of apoptotic cell death in melanoma cells as compared with normal keratinocytes using a non-thermal plasma torch. Cancer Biol Ther. 2012;13(13):1299–306. doi: 10.4161/cbt.21787.

8. Vandamme M, Robert E, Lerondel S, Sarron V, Ries D, Dozias S, et al. ROS implication in a new antitumor strategy based on non-thermal plasma. Int J Cancer. 2012;130(9):2185–94. doi: 10.1002/ijc.26252.

9. Guerrero-Preston R, Ogawa T, Uemura M, Shumulinsky G, Valle BL, Pirini F, et al. Cold atmospheric plasma treatment selectively targets head and neck squamous cell carcinoma cells. Int J Mol Med. 2014;34(4):941–6. doi: 10.3892/ijmm.2014.1849.

10. Hirst AM, Simms MS, Mann VM, Maitland NJ, O’Connell D, Frame FM. Low-temperature plasma treatment induces DNA damage leading to necrotic cell death in primary prostate epithelial cells. Br J Cancer. 2015;112(9):1536–45. doi: 10.1038/bjc.2015.113.

11. Adachi T, Tanaka H, Nonomura S, Hara H, Kondo S, Hori M. Plasma-activated medium induces A549 cell injury via a spiral apoptotic cascade involving the mitochondrial-nuclear network. Free Radic Biol Med. 2015;79:28–44. doi: 10.1016/j.freeradbiomed.2014.11.014.

12. Saito K, Asai T, Fujiwara K, Sahara J, Koguchi H, Fukuda N, et al. Tumor-selective mitochondrial network collapse induced by atmospheric gas plasma-activated medium. Oncotarget. 2016;7(15):19910–27. doi: 10.18632/oncotarget.7889.

13. Tokunaga T, Ando T, Suzuki-Karasaki M, Ito T, Onoe-Takahashi A, Ochiai T, et al. Plasma-stimulated medium kills TRAIL-resistant human malignant cells by promoting caspase-independent cell death via membrane potential and calcium dynamics modulation. Int J Oncol. 2018;52(3):697–708. doi: 10.3892/ijo.2018.4251.

14. Ito T, Ando T, Suzuki-Karasaki M, Tokunaga T, Yoshida Y, Ochiai T, et al. Cold PSM, but not TRAIL, triggers autophagic cell death: A therapeutic advantage of PSM over TRAIL. Int J Oncol. 2018;53(2):503–14. doi: 10.3892/ijo.2018.4413.

15. Tornin J, Mateu-Sanz M, Rodríguez A, Labay C, Rodríguez R, Canal C. Pyruvate Plays a Main Role in the Antitumoral Selectivity of Cold Atmospheric Plasma in Osteosarcoma. Sci Rep. 2019;9(1):10681. doi: 10.1038/s41598-019-47128-1.

16. Azzariti A, Iacobazzi RM, Di Fonte R, Porcelli L, Gristina R, Favia P, et al. Plasma-activated medium triggers cell death and the presentation of immune activating danger signals in melanoma and pancreatic cancer cells. Sci Rep. 2019;9(1):4099. doi: 10.1038/s41598-019-40637-z.

17. Ando T, Suzuki-Karasaki M, Ichikawa J, Ochiai T, Yoshida Y, Haro H, et al. Combined Anticancer Effect of Plasma-Activated Infusion and Salinomycin by Targeting Autophagy and Mitochondrial Morphology. Front Oncol. 2021;11:593127. doi: 10.3389/fonc.2021.593127.

18. Hollenbeck PJ, Saxton WM. The axonal transport of mitochondria. J Cell Sci. 2005;118(Pt 23):5411–9. doi: 10.1242/jcs.02745.

19. Cunniff B, McKenzie AJ, Heintz NH, Howe AK. AMPK activity regulates trafficking of mitochondria to the leading edge during cell migration and matrix invasion. Mol Biol Cell. 2016;27(17):2662–74. doi: 10.1091/mbc.E16-05-0286.

20. Landes T, Martinou JC. Mitochondrial outer membrane permeabilization during apoptosis: the role of mitochondrial fission. Biochim Biophys Acta. 2011;1813(4):540–5. doi: 10.1016/j.bbamcr.2011.01.021.

21. Elgass K, Pakay J, Ryan MT, Palmer CS. Recent advances into the understanding of mitochondrial fission. Biochim Biophys Acta. 2013;1833(1):150–61. doi: 10.1016/j.bbamcr.2012.05.002.

22. Twig G, Shirihai OS. The interplay between mitochondrial dynamics and mitophagy. Antioxid Redox Signal. 2011;14(10):1939–51. doi: 10.1089/ars.2010.3779.

23. Hoppins S, Lackner L, Nunnari J. The machines that divide and fuse mitochondria. Annu Rev Biochem. 2007;76:751–80. doi: 10.1146/annurev.biochem.76.071905.090048.

24. Soubannier V, McBride HM. Positioning mitochondrial plasticity within cellular signaling cascades. Biochim Biophys Acta. 2009;1793(1):154–70. doi: 10.1016/j.bbamcr.2008.07.008.

25. Pendin D, Filadi R, Pizzo P. The Concerted Action of Mitochondrial Dynamics and Positioning: New Characters in Cancer Onset and Progression. Front Oncol. 2017;7:102. doi: 10.3389/fonc.2017.00102.

26. Bell EL, Klimova TA, Eisenbart J, Schumacker PT, Chandel NS. Mitochondrial reactive oxygen species trigger hypoxia-inducible factor-dependent extension of the replicative life span during hypoxia. Mol Cell Biol. 2007;27(16):5737–45. doi: 10.1128/MCB.02265-06.

27. Desireddi JR, Farrow KN, Marks JD, Waypa GB, Schumacker PT. Hypoxia increases ROS signaling and cytosolic Ca(2+) in pulmonary artery smooth muscle cells of mouse lungs slices. Antioxid Redox Signal. 2010;12(5):595–602. doi: 10.1089/ars.2009.2862.

28. Semenza GL. Regulation of metabolism by hypoxia-inducible factor 1. Cold Spring Harb Symp Quant Biol. 2011;76:347–53. doi: 10.1101/sqb.2011.76.010678.

29. Al-Mehdi AB, Pastukh VM, Swiger BM, Reed DJ, Patel MR, Bardwell GC, et al. Perinuclear mitochondrial clustering creates an oxidant-rich nuclear domain required for hypoxia-induced transcription. Sci Signal. 2012;5(231):ra47. doi: 10.1126/scisignal.2002712.

30. Akita M, Suzuki-Karasaki M, Fujiwara K, Nakagawa C, Soma M, Yoshida Y, et al. Mitochondrial division inhibitor-1 induces mitochondrial hyperfusion and sensitizes human cancer cells to TRAIL-induced apoptosis. Int J Oncol. 2014;45(5):1901–12. doi: 10.3892/ijo.2014.2608.

31. Suzuki-Karasaki Y, Fujiwara K, Saito K, Suzuki-Karasaki M, Ochiai T, Soma M. Distinct effects of TRAIL on the mitochondrial network in human cancer cells and normal cells: role of plasma membrane depolarization. Oncotarget. 2015;6(25):21572–88. doi: 10.18632/oncotarget.4268.

32. Tanaka Y, Kanai Y, Okada Y, Nonaka S, Takeda S, Harada A, et al. Targeted disruption of mouse conventional kinesin heavy chain, kif5B, results in abnormal perinuclear clustering of mitochondria. Cell. 1998;93(7):1147–58. doi: 10.1016/s0092-8674(00)81459-2.

33. Pilling AD, Horiuchi D, Lively CM, Saxton WM. Kinesin-1 and Dynein are the primary motors for fast transport of mitochondria in Drosophila motor axons. Mol Biol Cell. 2006;17(4):2057–68. doi: 10.1091/mbc.e05-06-0526.

34. Lucken-Ardjomande Hasler S. Cholesterol, cardiolipin, and mitochondria permeabilisation. Anticancer Agents Med Chem. 2012;12(4):329–39. doi: 10.2174/187152012800228724.

35. Kaminskyy VO, Zhivotovsky B. Free radicals in cross talk between autophagy and apoptosis. Antioxid Redox Signal. 2014;21(1):86–102. doi: 10.1089/ars.2013.5746.

36. Ahmadpour ST, Mahéo K, Servais S, Brisson L, Dumas JF. Cardiolipin, the Mitochondrial Signature Lipid: Implication in Cancer. Int J Mol Sci. 2020;21(21). doi: 10.3390/ijms21218031.

37. Petit JM, Maftah A, Ratinaud MH, Julien R. 10N-nonyl acridine orange interacts with cardiolipin and allows the quantification of this phospholipid in isolated mitochondria. Eur J Biochem. 1992;209(1):267–73. doi: 10.1111/j.1432-1033.1992.tb17285.x.

38. Dixon SJ, Lemberg KM, Lamprecht MR, Skouta R, Zaitsev EM, Gleason CE, et al. Ferroptosis: an iron-dependent form of nonapoptotic cell death. Cell. 2012;149(5):1060–72. doi: 10.1016/j.cell.2012.03.042.

39. Miotto G, Rossetto M, Di Paolo ML, Orian L, Venerando R, Roveri A, et al. Insight into the mechanism of ferroptosis inhibition by ferrostatin-1. Redox Biol. 2020;28:101328. doi: 10.1016/j.redox.2019.101328.

40. Tochigi M, Inoue T, Suzuki-Karasaki M, Ochiai T, Ra C, Suzuki-Karasaki Y. Hydrogen peroxide induces cell death in human TRAIL-resistant melanoma through intracellular superoxide generation. Int J Oncol. 2013;42(3):863–72. doi: 10.3892/ijo.2013.1769.

41. Ohshima Y, Takata N, Suzuki-Karasaki M, Yoshida Y, Tokuhashi Y, Suzuki-Karasaki Y. Disrupting mitochondrial Ca2+ homeostasis causes tumor-selective TRAIL sensitization through mitochondrial network abnormalities. Int J Oncol. 2017;51(4):1146–58. doi: 10.3892/ijo.2017.4096.

42. Yoshikawa N, Liu W, Nakamura K, Yoshida K, Ikeda Y, Tanaka H, et al. Plasma-activated medium promotes autophagic cell death along with alteration of the mTOR pathway. Sci Rep. 2020;10(1):1614. doi: 10.1038/s41598-020-58667-3.

43. Gozuacik D, Kimchi A. Autophagy as a cell death and tumor suppressor mechanism. Oncogene. 2004;23(16):2891–906. doi: 10.1038/sj.onc.1207521.

44. Fulda S, Kögel D. Cell death by autophagy: emerging molecular mechanisms and implications for cancer therapy. Oncogene. 2015;34(40):5105–13. doi: 10.1038/onc.2014.458.

45. Gaschler MM, Hu F, Feng H, Linkermann A, Min W, Stockwell BR. Determination of the Subcellular Localization and Mechanism of Action of Ferrostatins in Suppressing Ferroptosis. ACS Chem Biol. 2018;13(4):1013–20. doi: 10.1021/acschembio.8b00199.

46. Quintana A, Schwarz EC, Schwindling C, Lipp P, Kaestner L, Hoth M. Sustained activity of calcium release-activated calcium channels requires translocation of mitochondria to the plasma membrane. J Biol Chem. 2006;281(52):40302–9. doi: 10.1074/jbc.M607896200.

47. Hajnóczky G, Saotome M, Csordás G, Weaver D, Yi M. Calcium signalling and mitochondrial motility. Novartis Found Symp. 2007;287:105–17; discussion 17-21. doi: 10.1002/9780470725207.ch8.

48. Niescier RF, Chang KT, Min KT. Miro, MCU, and calcium: bridging our understanding of mitochondrial movement in axons. Front Cell Neurosci. 2013;7:148. doi: 10.3389/fncel.2013.00148.

49. Madreiter-Sokolowski CT, Ramadani-Muja J, Ziomek G, Burgstaller S, Bischof H, Koshenov Z, et al. Tracking intra-and inter-organelle signaling of mitochondria. FEBS J. 2019;286(22):4378–401. doi: 10.1111/febs.15103.

50. Bravo R, Vicencio JM, Parra V, Troncoso R, Munoz JP, Bui M, et al. Increased ER-mitochondrial coupling promotes mitochondrial respiration and bioenergetics during early phases of ER stress. J Cell Sci. 2011;124(Pt 13):2143–52. doi: 10.1242/jcs.080762.

51. Mironov SL, Ivannikov MV, Johansson M. [Ca2+]i signaling between mitochondria and endoplasmic reticulum in neurons is regulated by microtubules. From mitochondrial permeability transition pore to Ca2+-induced Ca2+ release. J Biol Chem. 2005;280(1):715–21. doi: 10.1074/jbc.M409819200.

52. Debattisti V, Gerencser AA, Saotome M, Das S, Hajnóczky G. ROS Control Mitochondrial Motility through p38 and the Motor Adaptor Miro/Trak. Cell Rep. 2017;21(6):1667–80. doi: 10.1016/j.celrep.2017.10.060.

53. Alshaabi H, Shannon N, Gravelle R, Milczarek S, Messier T, Cunniff B. Miro1-mediated mitochondrial positioning supports subcellular redox status. Redox Biol. 2021;38:101818. doi: 10.1016/j.redox.2020.101818.

