## Supplementary for "Air plasma-activated medium exerts tumor-specific cytotoxicity via the oxidative stress-induced perinuclear mitochondrial clustering"

**Supplementary Figure S1**

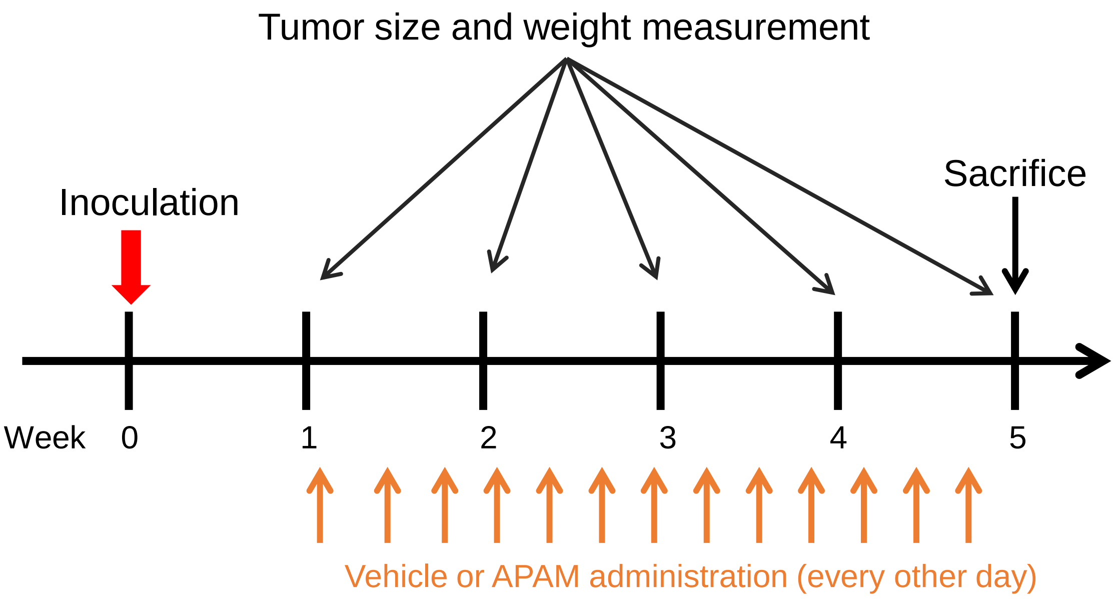

### Supplementary Figure S2

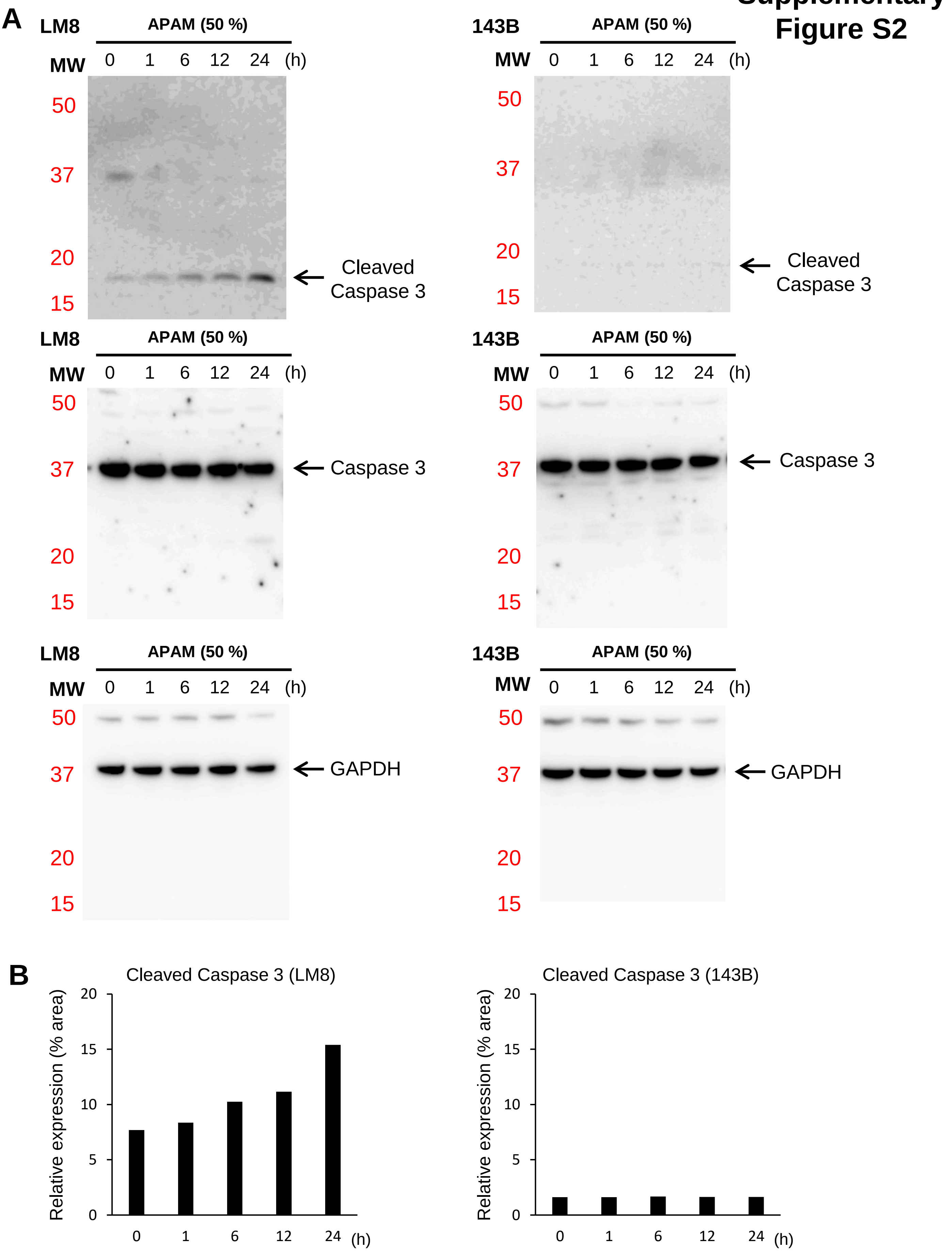

#### Supplementary Figure S3

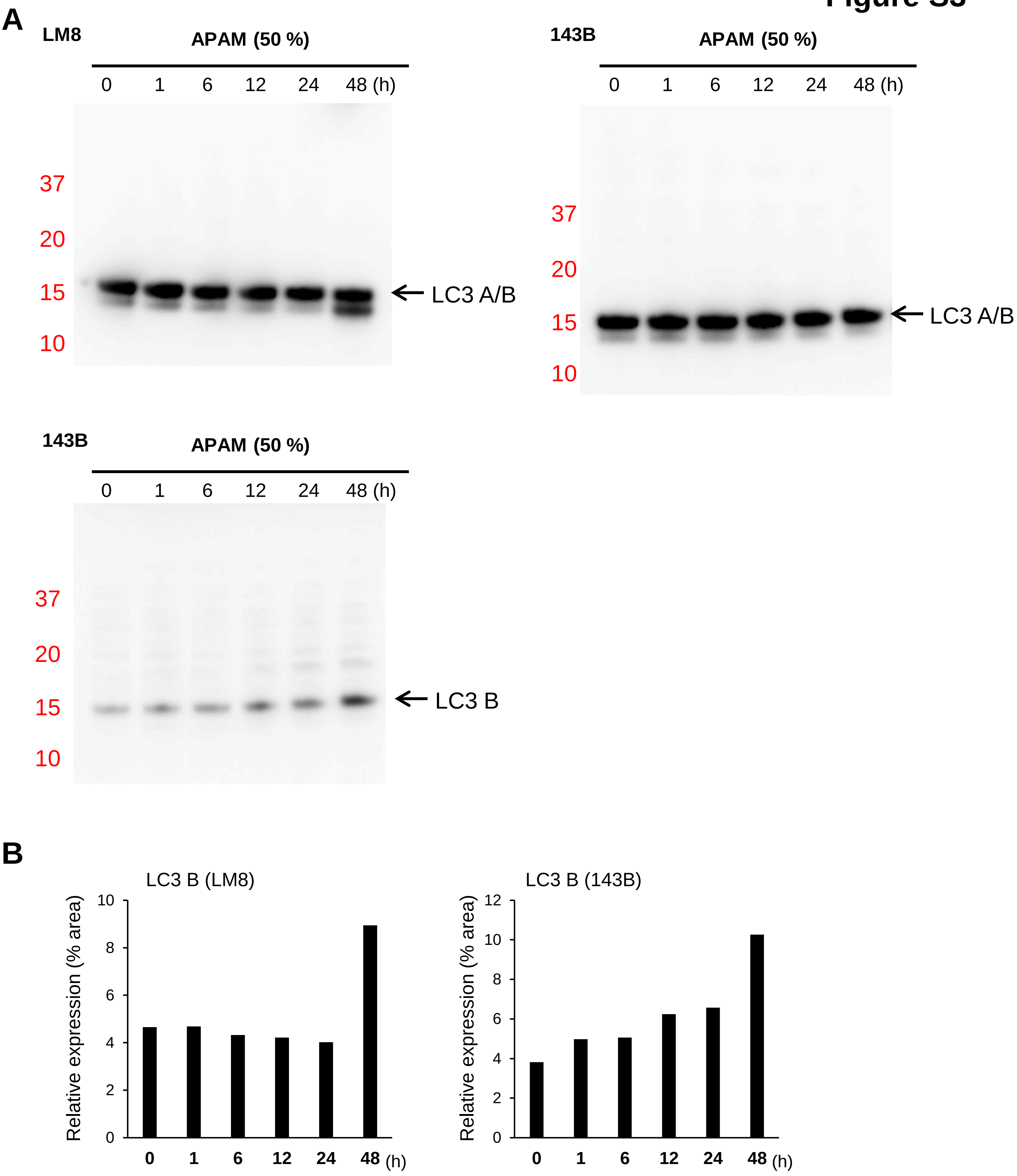

**Supplementary Figure S4**

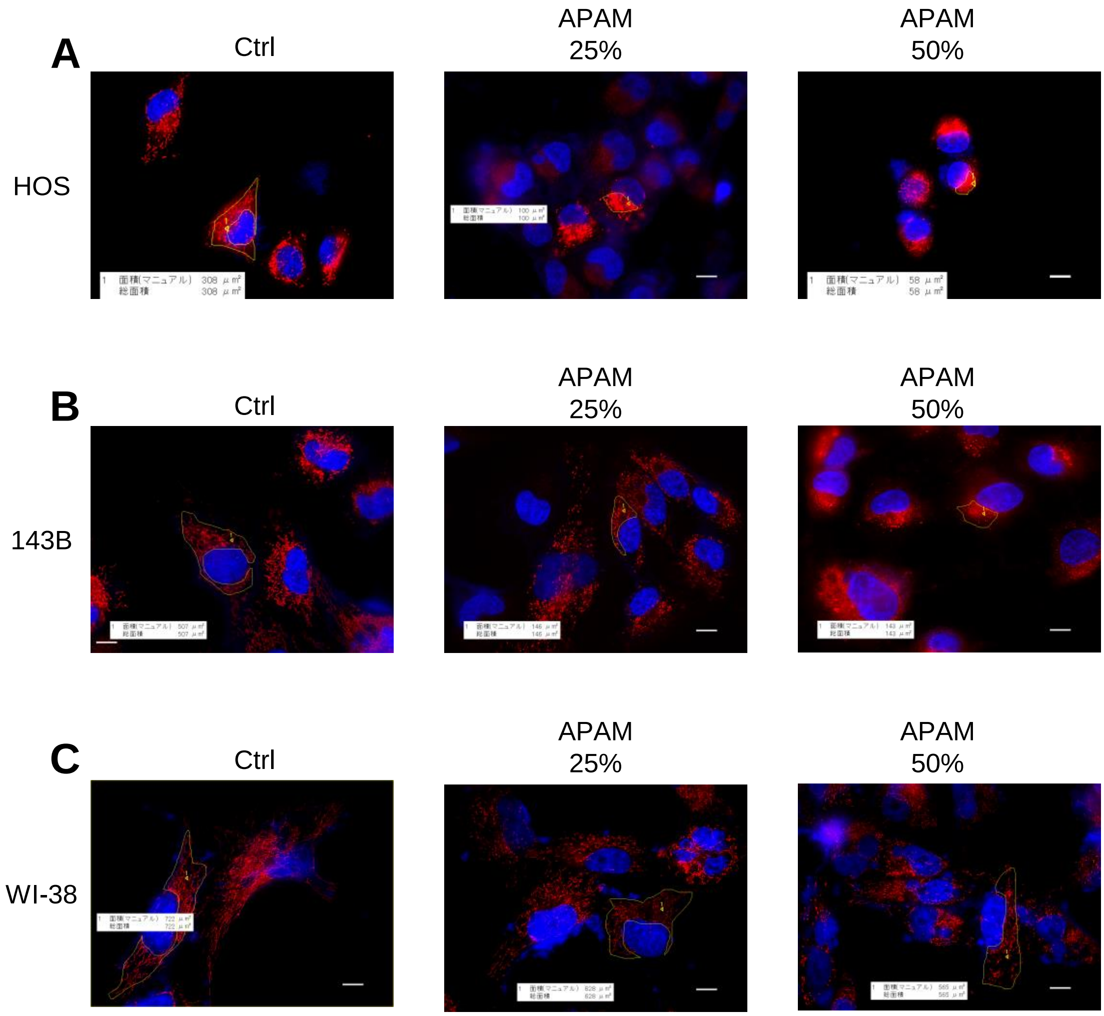
